# Xist ribonucleoproteins promote female sex-biased autoimmunity

**DOI:** 10.1101/2022.11.05.515306

**Authors:** Diana R. Dou, Yanding Zhao, Julia A. Belk, Yang Zhao, Kerriann M. Casey, Derek C. Chen, Rui Li, Bingfei Yu, Suhas Srinivasan, Brian T. Abe, Katerina Kraft, Ceke Hellström, Ronald Sjöberg, Sarah Chang, Allan Feng, Daniel W. Goldman, Ami A. Shah, Michelle Petri, Lorinda S. Chung, David F. Fiorentino, Emma K. Lundberg, Anton Wutz, Paul J. Utz, Howard Y. Chang

## Abstract

Autoimmune diseases disproportionately affect females more than males. The XX sex chromosome complement is strongly associated with susceptibility to autoimmunity. Xist long noncoding RNA (lncRNA) is expressed only in females to randomly inactivate one of the two X chromosomes to achieve gene dosage compensation. Here, we show that the Xist ribonucleoprotein (RNP) complex, comprised of numerous autoantigenic components, is an important driver of sex-biased autoimmunity. Inducible transgenic expression of a non-silencing form of *Xist* in male mice introduced Xist RNP complexes and sufficed to produce autoantibodies. Male SJL/J mice expressing transgenic Xist developed more severe multiorgan pathology in pristane-induced model of lupus than wild-type males. Xist expression in males reprogrammed T and B cell population and chromatin states to more resemble wild type females. Human patients with autoimmune diseases displayed significant autoantibodies to multiple components of XIST RNP. Thus, a sex-specific lncRNA scaffolds ubiquitous RNP components to drive sex-biased immunity.

**HIGHLIGHTS:**

- Transgenic mouse models inducibly express Xist in male animals.
- Xist expression in males induce autoantibodies and autoimmune pathology.
- Xist in males reprograms T and B cell populations to female-like patterns.
- Autoantibodies to Xist RNP characterize female-biased autoimmune diseases.

## INTRODUCTION

Autoimmune diseases are the third-most prevalent disease category, outpaced only by cancer and heart disease^1^.Four out of five patients with autoimmune diseases are female. For instance, in systemic lupus erythematosus (SLE), the ratio of patient sex is is 9:1 females to males; the ratio in Sjogren’s disease is 19:1 female to male patients^2,3^. Although hormones have been extensively studied^4^, the dosage of X chromosome appears to be a major driver of autoimmune risk irrespective of sex or hormonal status in humans and mice^5–8^. Patients with Klinefelter syndrome (XXY) are phenotypically males, have male hormonal pattern, but have an elevated risk of autoimmune disease equivalent to females. Specific X-linked genes, such as *TLR7,* that can escape X inactivation have been nominated as contributors to specific autoimmune diseases^5–8^. The genetic risk underlying autoimmune diseases from the second X chromosome in aggregate remain unresolved. In addition, identical twin studies have also shown varying degrees of autoimmune disease penetrance, suggesting a genetic disposition that is also reliant on environmental factors^9,10^. Hence, adjuvant triggers^11^ in addition to genetic predisposition may be initiators of autoimmune disease development.

Mammalian females have a XX genotype and males have a XY genotype. To make the gene expression output roughly equivalent between females and males, every cell in a female’s body epigenetically silences one of two X chromosomes via the action of the long noncoding RNA Xist. Xist is a ∼17 kb lncRNA (19 kb in human) that is transcribed only from the inactive X chromosome, and thus not expressed in males. Xist is critical for the establishment of X chromosome inactivation (XCI) spreading from the X-inactivation center and coating the entire inactive X in association with its protein partners. During XCI establishment in mouse embryonic stem cells, Xist associates with 81 unique binding proteins to form an RNP complex, 10 through direct RNA-protein interaction and others through indirect protein-protein interaction^12,13^. Xist is widely expressed in adult somatic tissues and associates additional tissue-specific proteins^14^. Several Xist binding proteins were previously noted to be autoantigens^12^. Studies in SLE patients and mice demonstrated that DNA-autoantibody and RNA-autoantigen immune complexes, such as Sm/RNP and U1A, activate the TLR7, TLR8 and TLR9 pathways of the innate immune system^15–17^. The XIST RNP, comprised of a lncRNA, bound RNA binding proteins, and tethered to pieces of genomic DNA, presents qualities resembling nucleic acid-autoantigen immune complexes.

In order to study the impact of the XIST RNP, independent of hormonal background, in autoimmune predilection, we utilized an inducible and non-silencing allele of *Xist* introduced into an autosome in the autoimmune-resistant C57BL/6J and autoimmune-prone SJL/J strain backgrounds. Inducing transgenic Xist RNP formation in male animals allowed the study of this female-specific lncRNA in a male background in the context of a chemically-induced SLE model. Both increased disease severity and elevated autoreactive lymphocyte pathway signatures were observed in the mouse models of pristane-induced SLE. Concurrently, we designed an antigen array to test autoimmune patient seroreactivity to XIST-associating proteins and detected significant reactivity towards multiple components of the XIST RNP. Altogether, our data point to a significant role for the *Xist* RNP as a driver for autoimmunity that may underly the sex-biased female preponderance for developing autoimmune diseases.

## RESULTS

### Xist RNPs as autoantigens in human disease

A defining feature of many autoimmune diseases is the development of antibodies against self-proteins, termed autoantibodies. Many autoantibodies are directed toward nuclear RNA binding proteins, and the nature and titer of such autoantibodies define the type and severity of autoimmune diseases in clinical practice. We and others have identified the constellation of RNA binding proteins associated with Xist RNA in several cell types^12–14^. Bibliomic analysis revealed that 30 proteins of Xist RNP constituents have been reported as the targets of autoantibodies (i.e., autoantigens) in one or more human diseases (**Supplementary Figure 1, Supplementary Table 1**). This observation stimulated the hypothesis that Xist RNP may promote female-biased autoimmunity.

### Developing the TetOP-DRepA-Xist transgenic mouse to model autoimmune diseases

To test Xist ribonucleoprotein as a potential trigger of autoimmunity, we developed a TetOP-**Δ**RepA-Xist transgenic mouse that enables inducible expression of *Xist* in male animals. Because Xist expression from an autosome silences the chromosome in cis and is often cell lethal, we chose to use **Δ**RepA-Xist, a truncation of Xist that removes the A-repeat (RepA) element required for gene silencing activity of Xist ^18^, but does not ablate chromosome coating or Xist RNP formation. Previous study indicated that 78 of 81 proteins in the Xist RNP associates with **Δ**RepA-Xist (Chu et al., 2015). Expression of the **Δ**RepA mutant Xist is controlled through the Tet-operon promoter (TetOP), and the transgenic cassette is inserted in the *Col1A1* locus on chromosome 11 (**Figure 1A**). Since Xist is expressed on only 1 of the 2 X chromosomes, we used mice heterozygous for TetOP-**Δ**RepA-Xist (denoted as tgXist onwards) in our studies. After only 2 weeks of doxycycline administration in heterozygous tgXist male mice, expression of *tgXist* was detectable through *Xist* qRT-PCR in multiple tissues and as single punctate foci in the nucleus reminiscent of Barr body as evidenced by RNA fluorescent in situ hybridization (FISH) (**Figure 1B,C**). We note that in the absence of induction, tgXist has low level of expression that is detectable by qRT-PCR and by FISH; upon induction, tgXist level increases ∼100-fold to approximate the level of endogenous Xist in female tissues (**Figure 1B-D**). tgXist did not reduce chromatin accessibility or RNA expression across chromosome 11 or locally at the locus of transgene insertion (**Supplementary Figure 2A-D**), consistent with the notion that **Δ**RepA-Xist is functionally null for gene silencing^19^. Backcrossing of the inducible tgXist transgene into common mouse strains enables the study of *Xist* in male animals in multiple autoimmune disease models.

**Figure 1:**
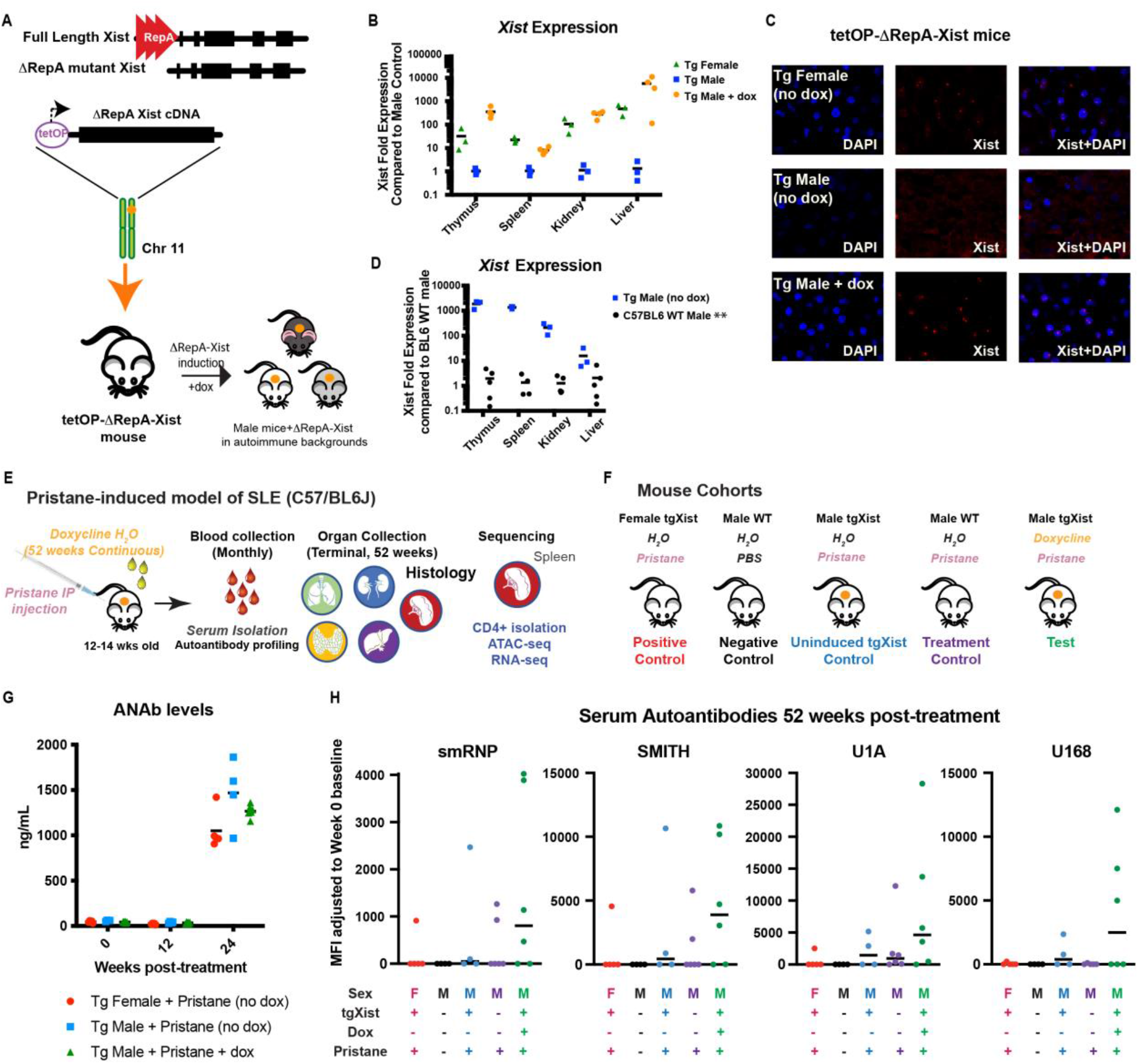
tgXist-mice inducibly express ΔrepA-Xist under doxycycline exposure. (A) Transgene construct of truncated ΔrepA Xist under tet-o-promoter in mice. (B) qRT-PCR and (C) FISH of Xist expression in tgXist-mice after 2 weeks of doxycycline administered in drinking water. (D) qRT-PCR comparison of basal tgXist levels in tgXist male mice without doxycycline exposure and WT male mice. For B and D, number of tgXist Female = 3, tgXist Male (no dox) = 3, tgXist Male + dox = 4, Wt Male = 5. (E) Schematic of tissue collection in pristane-induced SLE model in the C57/BL6J strain. (F) Cohorts in the induced SLE model. (G) ELISA of Serum ANAb levels, number of tgXist Female = 5, tgXist Male (no dox) = 4, tgXist Male + dox = 6. H) Median Fluorescence Intensity (MFI) of serum reactivity to autoantigens using a bead-based lupus antigen array after 52 weeks of treatment. From left to right tgXist Female control to tgXist Male test, number of animals in serum studies: n=5, n=4, n=4, n=6, n=6.

### Transgenic Xist promotes autoantibody production in males

To test our hypothesis that *Xist* RNPs creates increased susceptibility to autoimmune diseases, male and female C57BL/6J mice heterozygous for the ΔRepA-Xist transgene were injected with pristane to chemically induce systemic lupus erythematosus (SLE)^20–23^ and evaluated for disease 1 year post-treatment (**Figure 1E**). The pristane model of SLE has a well-documented female bias in disease penetrance and severity. Since C57BL/6J mouse^20,24^, the most common and widely used murine experimental background, are autoimmune resistant and male mice are not expected to develop pristane-induced SLE^20^, we first tested the effect of *tgXist* expressing on surrogate markers of autoimmunity.

The pristane-injected transgenic cohorts consisted of a female control untreated for doxycycline (female), male control untreated for doxycycline (tg male), and the test group of male mice continuously treated with doxycycline to induce *tgXist* expression (male+dox) (**Figure 1F**). Pristane-treated and untreated wild-type males of the same background were included as additional wild-type male controls. The rationale for each treatment cohort and comparison control is detailed in **Supplementary Table 2**. Measurement of anti-nuclear antibody (ANAb) production showed elevation of 24 weeks post-pristane injection in all transgenic cohorts tested, including the transgenic male mice not administered doxycycline (**Figure 1G**). Comparison of *tgXist* levels through qPCR between control transgenic male mice and wild-type C57BL/6 male mice suggests transgenic male mice already express a basal level of *tgXist* due to promoter leakage (**Figure 1D**). This level is low compared to the transgenic female control mice (**Figure 1B**) but may be sufficient to affect changes in autoimmunity. Since C57BL/6J mice are autoimmune-resistant, 52 weeks post-induction was the terminal timepoint for pristane-induced SLE^20^. However, the resulting advanced age of the mice combined with their natural autoimmune resistance made detection of disease-specific pathophysiological changes difficult to determine (data not shown).

Despite the lack of significant physiological SLE in any mice (including female controls), we detected elevated presence of autoantibodies to several SLE-associated antigens, all of which are RNA-binding proteins (snRNP, Smith proteins B/B’, U1-A, and U1-68) 52-weeks post-pristane treatment with the highest median elevation in the tgXist-induced male cohort. Since elevated autoantibody levels were detected across all three transgenic cohorts (female, male, and male+dox), we also injected wild-type C57BL/6J male mice with pristane (WT Male) or a mock injection of PBS (WT Male +PBS) to test for pristane effects on a wild-type background without basal levels of *tgXist*. While some elevated autoantibodies were also detected in one of four wildtype pristane-injected male mice, it does not approach the higher frequency and level of elevation observed in the tgXist-induced male group (**Figure 1H**).

### Transgenic Xist drives males to female-like changes in T cell profiles

While pristane-induced SLE in C57BL/6J mice did not physically manifest disease (data not shown), splenic CD4^+^ T cells (a critically important cell type for balancing and driving autoimmunity^25–27^) from *tgXist*-expressing male mice correlated closer to females than to control (tgXist-non-induced/wild-type) males at several levels. At the terminal timepoint of 52 weeks post-treatment, changes in the transcriptional regulation and gene expression of CD4+ splenic T cells were assessed using ATAC-seq and RNA-seq of multiple animals per cohort (**Figure 2A**). Global comparison of ATAC-seq differential peaks showed several differences between non-tgXist/Xist-expressing animals compared to tgXist-expressing males and wild-type Xist-expressing females, but very few differential peaks between females compared to males expressing tgXist (**Figure 2B**). PCA comparison of female mice, male mice, and male tgXist-expressing mice (male+dox) showed a separation of mice most highly correlated with sex (**Figure 2C**). Two of six tgXist-expressing male mice showed female-like skewing (circled in **Figure 2C**), in keeping with the expected female penetrance percentage and subsequent analysis focused on these two mice to determine the epigenetic and genomic changes contributing to the shift from male-like to female-like.

**Figure 2:**
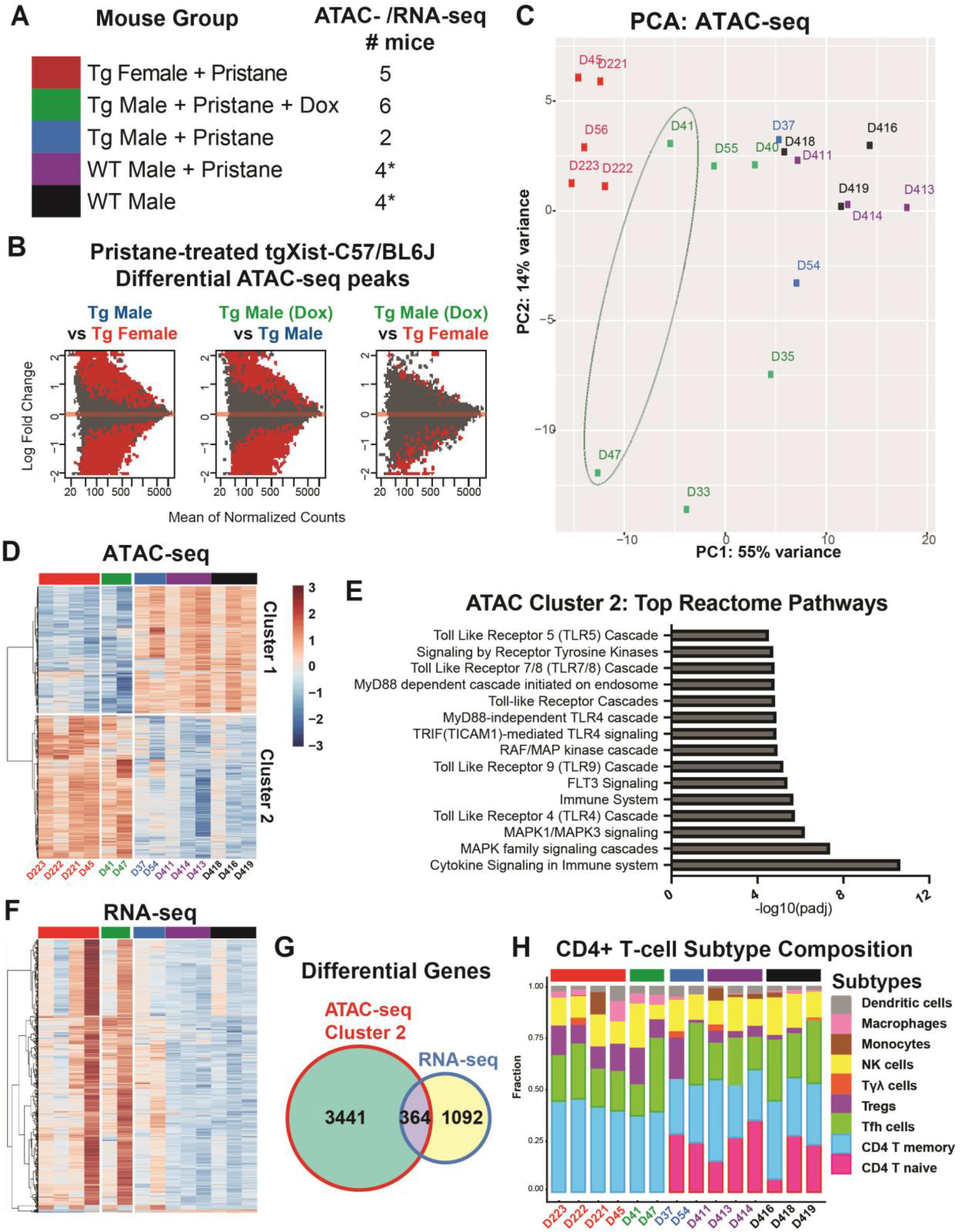
Bulk ATAC- and RNA-sequencing of splenic CD4+ T-cells from C57/BL6J mice reveal closer correlation between tgXist- and Xist-expressing mice in the pristane-induced SLE model. (A) Table of treatment cohorts: colors correspond to each treatment cohort and numbers indicate the number of ATAC- and RNA-seq samples. *Due to library quality, only 3 RNA-seq libraries were used. (B) MA plots comparing differential regions of genomic accessibility in transgenic cohorts. (C) Poly Component Analysis (PCA) plots of ATAC-seq libraries from all mice in the study. Heatmap of (D) ATAC-seq Z-scores and (F) RNA-seq differential gene expression from representative mice from each treatment cohort. (E) Top 15 differential reactomes associated with cluster 2 genomic regions. (G) Metrics of differential genes from cluster 2 ATAC-seq and RNA-seq datasets. (H) CIBERSORT prediction of T-cell subset composition from CD4+ T-cells RNA-seq gene expression libraries. (D)-(H): number of tgXist Female+pristane = 4, tgXist Male+pristane = 2, tgXist Male+dox+pristane = 2, Wt Male+pristane= 3, Wt Male mock treatment=3.

ATAC-seq revealed distinct clusters of accessibility corresponding to tgXist/Xist-expressing and non-expressing mice (**Figure 2D**). tgXist-induced and pristane-injected males displayed similar chromatin accessibility to pristane-injected female positive control mice, with higher accessibility in cluster 2, and were distinct from male negative control groups (*tgXist* non-induced and wild-type) that displayed higher accessibility in cluster 1 (**Figure 2D**). Interestingly, the top Reactome categories in Cluster 2, associated with *tgXist*-expressing males and Xist-expressing females, displayed high TLR pathway signatures (**Figure 2E**) not present in Cluster 1 (**Supplementary Figure 3D**). Notably, *Tlr9*, a pathogen sensor in the innate immune pathway that is highly active in SLE, is significantly more accessible in females and tgXist-induced males (**Supplementary Figure 3F**). RNA-seq profiles of gene expression displayed a similar trend and clustering grouped by tgXist/Xist expression with some variability within cohorts (**Figure 2F**), suggesting that ATAC-seq provides a more consistent profile of cells in transition. Comparison of dysregulated genes from ATAC- and RNA-seq revealed an overlap of 364 genes (**Figure 2G**). CIBERSORT^28^ deconvolution of gene expression signatures also identified a clear segregation of *Xist*-expressing and non-expressing mice. In particular, *tgXist or Xist-*expressing males and wild type females, respectively, contained more CD4 memory T-cells while control males expressed greater proportions of naïve T-cells (**Figure 2H**).

### Xist expression in males promotes multi-organ autoimmune pathology

We next tested the effect of tgXist expression in the autoimmune-prone SJL/J mouse background, a widely used strain in multiple autoimmune disease models, including pristane-induced SLE^21,29,30^. Pristane-induced SLE in SJLJ/J mice exhibit many characteristics of human SLE (i.e., autoantibody development, TLR7 upregulation, multi-organ involvement) and demonstrate a strong female bias (i.e., XX over XY genotypes)^10,21^. In sex-matched pristane-induced SLE studies, SJL/J female mice display earlier mortality, more severe nephritis, higher levels of autoantibodies, and are 3.4x more likely to die than male mice ^21^. This disease model is also more reflective of most human patients than spontaneous SLE models that are restricted to specific genetic mutations^31–34^.

We administered a one-time 0.5 mL intraperitoneal injection of pristane to 8-10 weeks old mice to induce SLE in *tgXist* and wild-type (WT) mice and continuous ingestion of doxycycline was supplied in drinking water to activate *tgXist* in selected mice (**Figure 3A**). The experimental cohorts include the wild-type female pristane-treated positive control, negative controls of mock-injected *tgXist*- and WT males, the WT male treatment control, and the test group of *tgXist*-expressing pristane-treated males (**Figure 3B, Supplementary Table 2**). Since previous literature determined that WT SJL/J females display mortality as early as 16 weeks post-pristane injection^21,21^, we selected the terminal timepoint of 16 weeks post-injection to avoid premature loss of mice.

**Figure 3:**
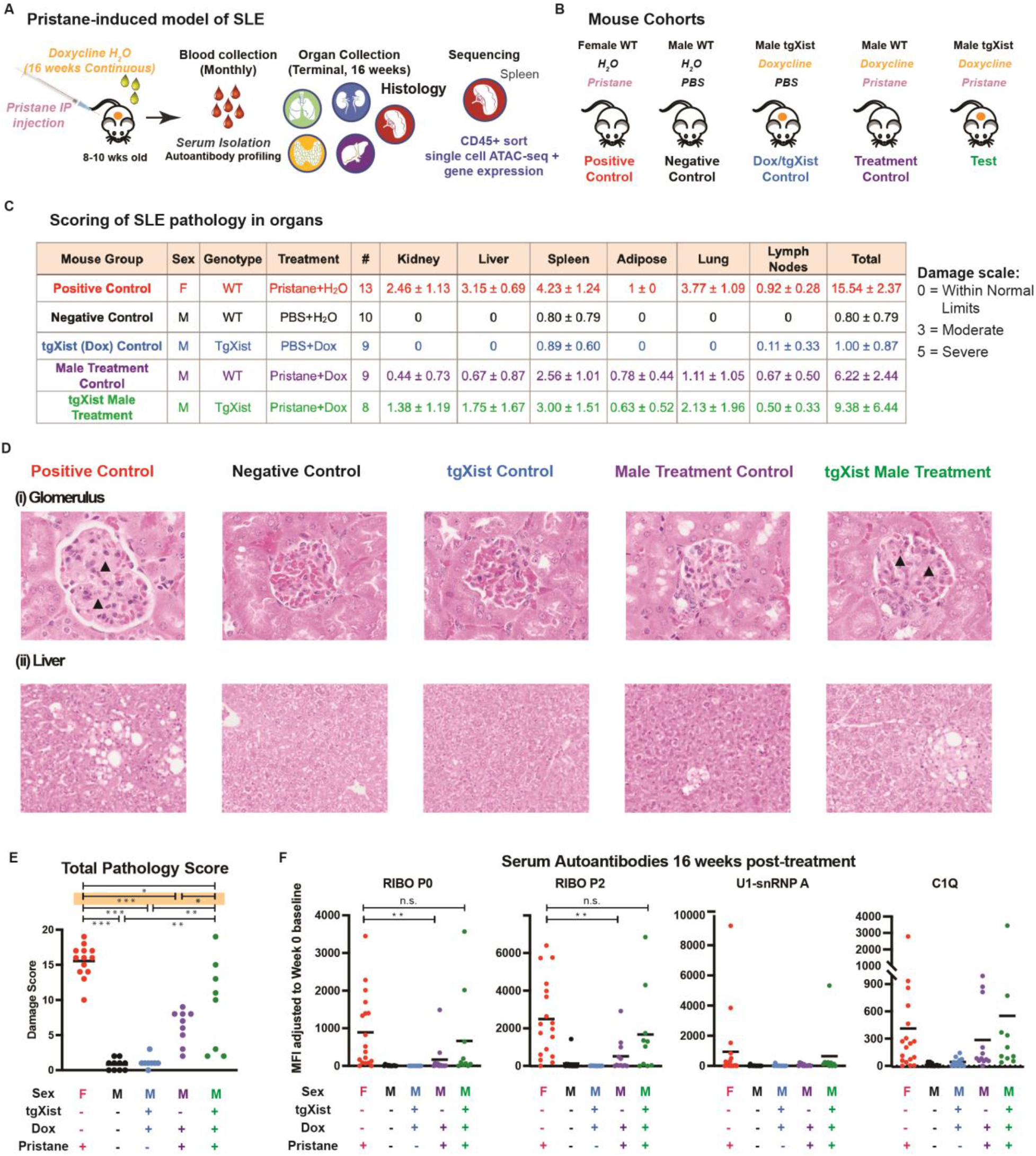
Increased phenotypic changes in pathophysiology and autoantigen levels in tgXist- and Xist-expressing mice in the SJL/J strain in the pristane-induced SLE model. (A) Schematic of pristane-induced SLE and strategy for histopathology, serum and sequencing analysis in the SJL/J strain. (B) Treatment cohorts in the SJL/J strain. (C) Table of the mean severity of damage across multiple tissue sites and mice. (D) Representative H&E images of (i) glomeruli (kidney) and (ii) liver sections from each treatment cohort. Arrows demarcate mesangial thickening. (E) Graph of total sum of pathophysiology damage scores. Significance calculated using the Fisher’s Exact Test, * indicates p < 0.05, ** p < 0.01, *** p < 0.001. (F) Median Fluorescence Intensity (MFI) of serum reactivity to autoantigens using a bead-based lupus antigen array after 16 weeks of treatment. Significance calculated using the unpaired Wilcoxon Rank Sum Test, * indicates p < 0.05, ** p < 0.01, *** p < 0.001, number of mouse serum samples, listed from left to right: wild-type Female + pristane =18, wild-type Male negative control=13, tgXist Male+dox (tgXist) control=11, wild-type Male treatment =12, tgXist Male treatment=10.

Since SLE is a systemic disease, we used H&E staining to assess the pathology of multiple affected organs at the terminal collection time (**Figure 3C,D**). In all pristane-treated cohorts, pristane injection caused lipogranulomas in adipose tissues, lymph node hyperplasia and medullary plasmocytosis, as well as varying degrees of extramedullary hematopoiesis and lymphoid expansion in the spleen. However, pristane injection coupled with *tgXist* expression in males or *Xist* in WT female resulted in greater incidence and severity of glomerulonephritis (kidney), hepatic lipogranulomas (liver), and pulmonary hemorrhage and lymphohistiocytic alveolitis (lung), which is reflective of disease damage in the kidney, liver, and lungs observed in severe SLE patients^35^ (**Figure 3C**). Manifestation of mild signs of early SLE in some pristane-injected male controls was not unexpected because WT SJL/J males are expected to develop milder disease and display later mortality at 24 weeks post-injection^21^.

We determined that the total systemic damage across all assessed organs significantly increased in pristane-treated *tgXist*-expressing males compared to pristane-treated WT males as well as untreated male controls (*p* < 0.05, Fisher’s Exact Test, **Figure 3E**). Comparatively, there was no significant difference in total damage scores between pristane-treated WT males and the untreated male controls. Notably, the greatest statistical difference was preserved in the comparison between the female positive controls and the pristane-treated WT males as it was between the positive female and negative male controls (**Figure 3E**, highlighted in orange).

Sera collected at treatment start (day 0, baseline) and terminally (16 weeks post-treatment) were assessed for reactivity to SLE antigens using the Luminex bead-based antigen array^36^. Both the female (WT *Xist*) and *tgXist* males treated with pristane demonstrated higher reactivity levels to SLE antigens, including the subunits of the ribosomal P antigen (Ribo P0, Ribo P2), U1-snRNP, and C1q complexes (**Figure 3F**). Notably, while WT males developed significantly lower Ribo P0 and P2 autoantibodies compared to WT females in the pristane-induced SLE treatment, there were no significant differences between the tgXist-induced, pristane-treated males and wild-type females while (**Figure 3F**). Based on these studies, similar to female SJL/J mice, *tgXist-*expressing male mice experience more severe disease than their wild-type male counterparts in the pristane-induced SLE model.

### Single-cell ATAC analysis reveals distinct cell type clustering and consistent reformatting of the chromatin landscape in pristane-treated SJL/J animals

To gain insight to the genes and cellular processes underpinning the heightened autoimmunity in tgXist and Xist-expressing mice, we created single cell ATAC+ gene expression libraries of CD45+ (pan-hematopoietic) sorted splenic cells from pristane-injected mice in the SJL/J strain. A key advantage of the single cell multiomics approach is the ability to interrogate all the hematopoietic lineages within the spleen in a single assay. While our bulk ATAC- and RNA-sequencing in the C57BL/6J strain was restricted to CD4+ T-cells, the single cell libraries encompass the whole CD45+ splenic population.

Using the total lowest observed pathology score in female mice (10) as the cutoff (**Figure 3B,D**), we divided the pristane-induced tgXist-expressing males into disease high (total pathology score ≧ 10) and low (<10) disease groups. Sequencing libraries were made from representative mice selected from the tgXist male high disease (tgM high, n=4), tgXist male low disease (tgM low, n=4), and two pristane-injected control groups: wild-type females (Wt F, n=4) and wild-type males (Wt M, n=3). Due to high mitochondrial/ribosomal RNA content, 2 of the wild-type females were excluded from the gene expression analysis. To maintain consistency, we included only cells with both ATAC and gene expression reads.

The single cell ATAC UMAP formed 19 clusters (**Supplementary Figure 3A**) and were assigned cell type identities using imputation of key marker genes (**Figure 4A, Supplementary Figure 4C**). The four pristane-treated mouse group were generally evenly distributed across all clusters, although tgM high mice contained a higher fraction of cells within the CD4 Tcell cluster (**Figure 4B**). TgM high cells also formed a visually separate CD4_Tcell and Bcell_1 group on the UMAP (**Figure 4A, 4C**), but comparison of distinguishing markers did not show prominent differences from the adjacent CD4_Tcell and Bcell_1 populations. As expected, the tgXist male mice and female mice all showed peaks in the first *Xist* exon region (the *Xist* transcription start site is absent in the *Xist* transgene) that are lacking in the wild-type control males **(Figure 4D**).

**Figure 4:**
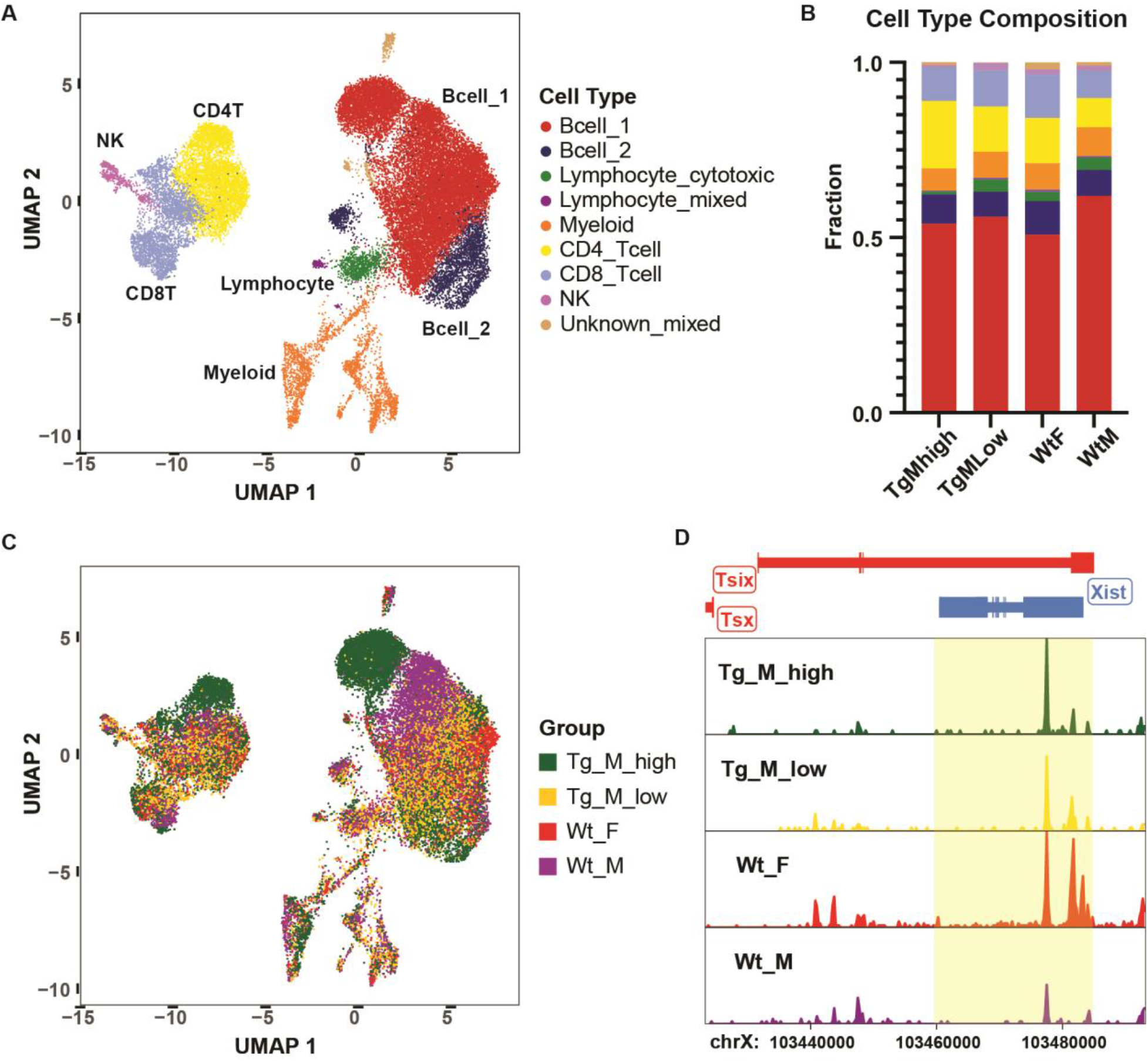
Single cell ATAC comparisons of splenic CD45+ hematopoietic cells from pristane-treated mice in the SJL/J strain. (A) UMAP of cell type cluster identities and (B) corresponding cell type composition within each pristane-treated mouse group. (C) UMAP clustering and (D) *Xist* peak tracks from pristane-treated mouse groups (features FDR ≦ 0.1 and Log2FC ≧ 0.5). Pristane-treated mouse groups shown: tgXist male disease high (n=4), tgXist male disease low (n=4), wild-type male (n=3), and wild-type female (n=2).

Interestingly, Wt F and tgM high cells appear to overlap in the main Bcell_2 cluster (**Figure 4A, 4C**), which colocalizes *Cd19* with the atypical B-cell marker *Zeb2* (**Supplementary Figure 4C**). However, pairwise comparisons between the mouse groups did not identify significant differences in the chromatin landscape within the main lymphocyte clusters even when comparing the most distinct positive disease control (Wt F) to the males with the lowest disease scores (Wt M and tgM low) (**Supplementary Figure 4D**). Aside from a higher CD4T cell fraction in the tgM high mice and atypical B cell activation markers, there were few noticeable distinguishing features in the single cell ATAC comparisons between the pristane-treated mouse groups, suggestive that all of the pristane-treated mice in the autoimmune-prone SJL/J model may have already undergone epigenetic remodeling preceding physiological disease onset at the time of investigation (16 weeks post-injection).

### Single-cell gene expression analysis reveals elevation of atypical B-cells and suppression of T-cell modulators in diseased *tgXist* animals

We next interrogated the pristane-treated SJL/J groups by single cell gene expression, which may reflect cell states with greater immediacy. The single cell gene expression UMAP formed 12 distinct clusters and were segregated clearly into B-cell, T-cell, NK cell, myeloid and erythroid lineages based on key transcription factors and markers (**Figure 5A, Supplementary Figure 5B**). The cellular type identities assigned to cells in the gene expression clusters closely matched the ATAC cell type identities (**Supplementary Figure 4B**), further validating the consistency of the assigned cellular identities in both the ATAC and gene expression datasets.

**Figure 5:**
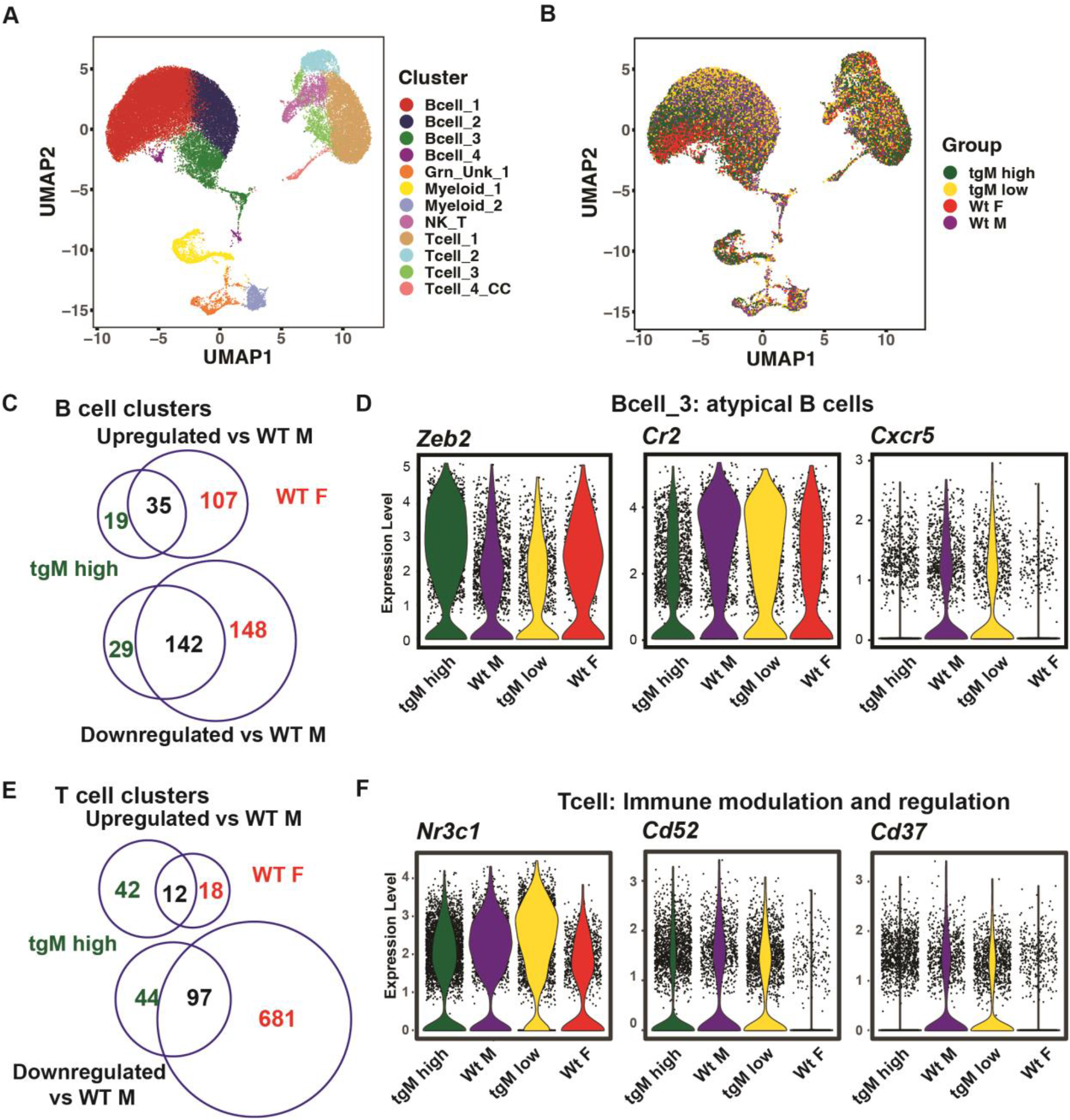
Single cell gene expression comparisons of splenic CD45+ hematopoietic cells from pristane-treated mice in the SJL/J strain. (A) Cell type cluster identities and (B) pristane-treated mouse groups displayed on the single cell gene expression clustering UMAP. (C) Metrics of differentially expressed B-cell cluster genes of tgXist Male high disease and wild-type female compared to wild-type males. (D) Representative gene expression plots of atypical B cell genes from the Bcell_3 cluster and (E) Metrics of differentially expressed T cell cluster genes of tgXist Male high disease and wild-type female compared to wild-type males. (F) T-cell cluster highlighting significant differentially expressed genes. Pristane-treated mouse groups shown: tgXist male disease high (n=4), tgXist male disease low (n=4), wild-type male (n=3), and wild-type female (n=2). Significance was calculated using the Wilcoxon Rank Sum test.

Within the B cell cluster, the tgXist male high disease group overlay with the female wild-type mice, distinct from the wildtype male and the relatively unaffected tgXist male low disease animals (**Figure 5B, Supplementary Figure 6A-D**). Since B cells produce the majority of autoantibodies and proinflammatory cytokines that characterize autoimmune pathogenesis^37,38^, the high correlation of B-cells, specifically, of the tgXist-diseased males with those from positive disease control females provides another layer of support for the hypothesis that Xist complexes mediate an environment of higher autoimmunity. The tgXist low disease and wild-type males grouped closely together and most of the differences between the tgXist high disease and wild-type female mice lay in the sex chromosomes or ribosomal protein genes. Inspection of the differentially expressed genes comparing the disease-affected tgXist male and female mice with the wild-type males (**Supplementary Figure 3C)** revealed that the majority of both downregulated and upregulated genes were shared between the first two groups (**Figure 5C**). Particularly significant genes included the upregulation of the atypical B-cell marker, *Zeb2* (tgM high vs WT M, *p* = 3.08E-69, WT F vs WT M, *p* = 1.17E-35) and *CD22* (tgM high vs WT M, *p* = 7.63E-46, WT F vs WT M *p* = 2.59E-36), a receptor associated with pathogenic B cells^39^ critical for B-cell proliferation and B cell receptor signaling (**Supplementary Figure 5D**). Concurrently, *Siglec-g*, encoding a receptor associated with promoting B-cell self-tolerance, deficiency of which is associated with increased B-1 cells and multiple autoimmune diseases^40–42^, was significantly downregulated (tgM high vs WT M, *p* = 1.73E-101, WT F vs WT M, p = 1.01E-43) (**Supplementary Figure 5D**). Also downregulated were complement receptor 2 (*Cr2*, tgM high vs WT M, *p* = 1.30E-153, WT F vs WT M *p* =3.70E-44) and the paralog to its alternatively spliced form, *Cr1l* (tgM high vs WT M, *p* = 4.41E-33, WT F vs WT M, *p* = 1.35E-33), both of which are important for suppressing autoimmunity^43,44^.

Within the B cell clusters, WT F and tgM high uniquely overlapped in a region of Bcell_3 not populated by either WT M or tgM low disease groups (**Supplementary Figure 6A-D**). Cluster Bcell_3 contained the highest correlation of gene expression signatures of atypical B-cell markers^45^ (**Supplementary Figure 6E**, highlighted in orange), including upregulation of *Cd19*, *Ms4a1* (gene encoding Cd20), *Zeb2*, and *Fcrl5,* a defining marker of atypical memory B-cells in both mice and humans^46^, as well as downregulation of *Cr2* (Cd21), *Cd27*, and *Cxcr5*^47,48^. Furthermore, Bcell_3 cluster cells from the gene expression data matched solely to the independently clustered multiome ATAC UMAP Bcell_2 cell clusters (**Supplementary Figure 3B**), the ATAC clusters imputed to have atypical B-cell characteristics (**Supplementary Figure 3C, *Zeb2* imputation**). Closer examination of the gene expression Bcell_3 cluster showed significant elevation of *Zeb2* (tgM high vs WT M, *p* = 6.12E-74, WT F vs WT M, *p* = 8.53E-9) and downregulation of *Cr2* (tgM high vs WT M, *p* = 1.75E-89, WT F vs WT M, *p* =3.03E-14) and *Cxcr5* (tgM high vs WT M, *p* = 5.65E-23, WT F vs WT M, *p* = 2.61E-18) in high disease compared to relatively unaffected mouse groups (representative plots, **Figure 5D**). CD21^-^CD27^-^ double negative are typical distinguishers of atypical B cells while loss of CXCR5 in CD21^-^ effector B cells is a hallmark of atypical B cells in SLE pathogenesis in patients^47,48^.

While the UMAP within the T cell clusters was less distinctly demarcated than the B-cell cluster (**Figure 5B, Supplementary Figure 5E**), the overall pattern was clear in the shared downregulated programs (**Figure 5E**). Multiple key T cell regulation and self-tolerance genes were downregulated in the tgXist male high disease and female cohorts compared to the wild-type males. Among the significantly downregulated genes were the glucocorticoid receptor *Nr3c1* (tgM high vs WT M, *p* = 1.37E-33, WT F vs WT M, *p* = 1.33E-63) involved in Treg modulation of inflammation^49^, *Cd37* (tgM high vs WT M, *p* = 4.60E-12, WT F vs WT M, *p* = 1.81E-10) which regulates proliferation^50^ and complement-mediated apoptosis of autoreactive T-cells^50^, the CD4 T-cell immune regulator *Cd52*^51^ (tgM high vs WT M, *p* = 7.24E-13, WT F vs WT M, *p* = 2.22E-72), and invariant chain *Cd74* involved in training antigen immunity in T cells^52^ (tgM high vs WT M, *p* = 8.15E-37, WT F vs WT M *p* = 2.27E-96) (representative plots, **Figure 5F**).

Our multiome sequencing identified clusters suggestive of atypical B cells overlapping closely in both ATAC and gene expression. At the single cell gene expression level, the severity of disease in tgXist-/Xist-expressing animals may be driven by increased atypical B cell activity and decreased immune modulatory programs in both B and T cells. Combined with the heightened pathophysiology scores and increased autoantigen levels, the gene expression data corroborates a role for *Xist* RNPs in the development of increased and/or more severe autoimmunity in the pristane-induced SLE model.

### Autoimmune patients display multiple autoantibodies to the XIST RNP

Although mouse models provide critical insights into biological mechanisms, there remain limitations in translating discoveries in mice to humans. In order to test whether the XIST RNP is immunogenic in humans, we obtained de-identified sera from patients with dermatomyositis (DM), scleroderma (SSc) and SLE to test for reactivity to XIST complex proteins^12,14^. We used protein fragments produced by the Human Protein Atlas for 128 of XIST-associated proteins of interest (selection described in Methods) and 52 for control proteins used as clinical markers for DM, SSc and SLE. Three of the clinical protein antigens overlapped with the XIST ChIRP-MS lists (SSB, SNRPD2, SRP14). When possible, multiple fragments spanning different regions of each protein were used (**Figure 6A**).

**Figure 6:**
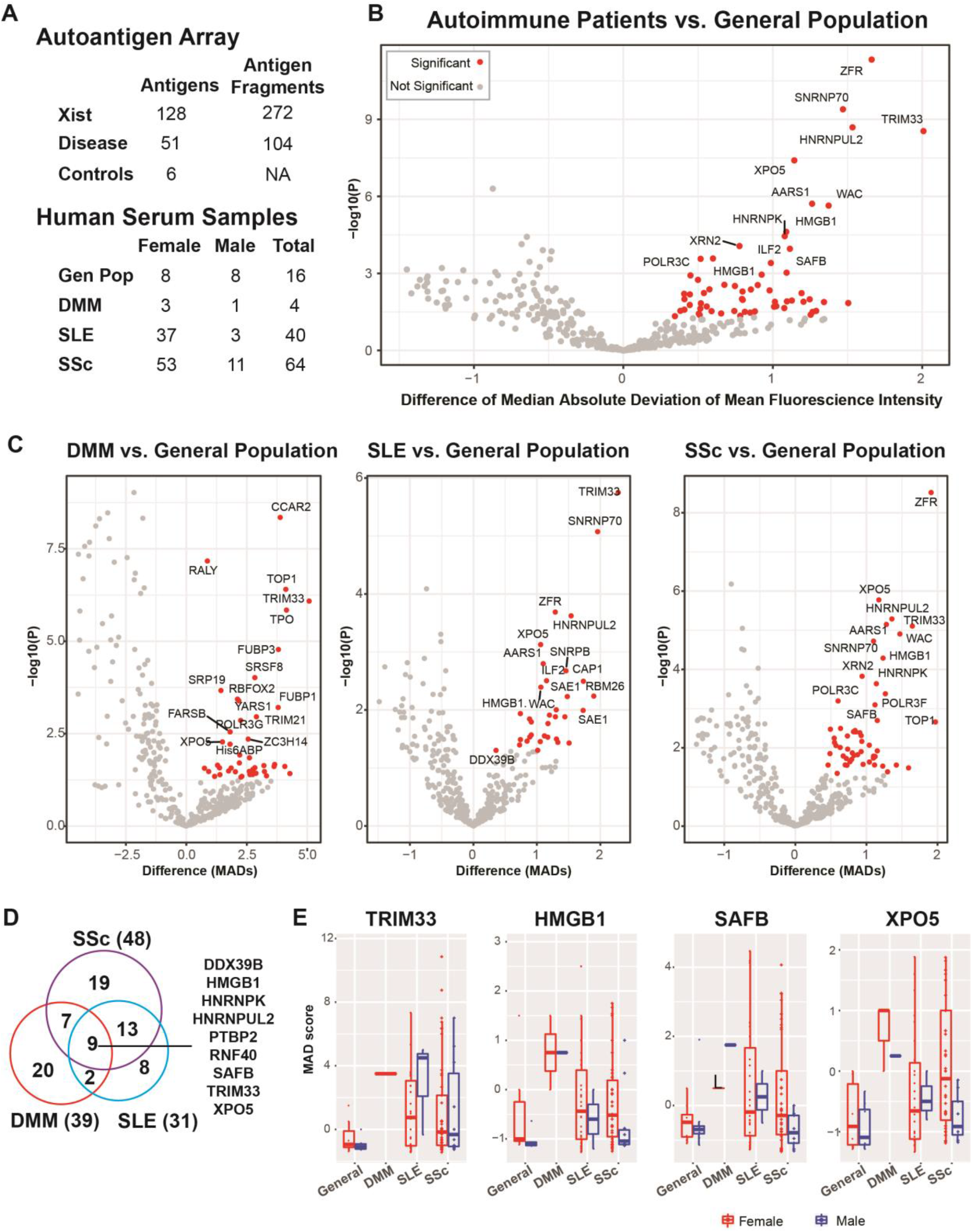
Autoimmune disease individuals experience increased serum reactivity towards antigens of the XIST RNP. (A) Table of the XIST complex-associated antigen array design showing the number of total unique proteins and corresponding protein fragments counts drawn from XIST ChIRP-MS, clinical disease panel, and control protein lists; and sample numbers of autoimmune patient and general population serum. (B) Volcano plot of serum reactivity of all autoimmune patients and (C) sera grouped by DM, SSc, and SLE patients compared to general population baseline of serum from blood donors. Significant differentially reactive antigens, defined as p_adj_ < 0.05 and MAD difference > 0, labeled in red. Significance calculated using the Student’s T-test. (D) Metrics of unique antigens from the array with significant elevated serum activity in autoimmune patients compared to the general population. (E) Serum reactivity (MAD) plots of representative antigens significantly reactive in all three autoimmune patient cohorts grouped by disease type and colored by sex (red=female, blue=male).

As a general population control, we obtained serum from anonymous donations to the Stanford Blood Center. However, since the mean and median age of these donors were 58 and 61 years at the time of donation, these donors may express some autoreactivity due to advanced age. Nevertheless, autoimmune patients were significantly more reactive to 55 proteins, 16 of which were disease markers, 1 of which overlapped with XIST RNP (SSB), and the remaining 39 antigens from the XIST RNP list (**Figure 6B, Supplementary Table 5**). Distinct reactive antigens for each autoimmune disease arose when grouped by disease (**Figure 6C**) and 9 antigens were shared among all 3 diseases (**Figure 6D, Supplementary Table 5**). Of these, TRIM33, or TIF1-γ, is a clinical marker for autoimmune disease (DM)^53^. The other 8 were XIST RNP components, several of which have recently discovered roles in autoimmunity. HMGB1 was recently identified as an autoantigen in SLE/Sjögren that may be another SS-protein in the SS-A/SS-B family^54,55^, HNRNPK is an autoantigen in a subset of Raynaud’s disease^56,57^ and aplastic anemia, SAFB is a novel autoantigen in connective tissues detected in interstitial lung disease, and XPO5 is in the SSB/La processing pathway^58^.

In sum, the 3 disease groups were significantly reactive to 79 unique proteins in the array compared to the general population control. Of these 79 proteins, 27 were disease controls, and 53 were associated with the XIST RNP (SSB is also a disease control marker). 37 of the 55 XIST-associated proteins were part of the group of 118 high confidence published XIST RNP complex proteins^14^. Of these 37 high confidence XIST complex proteins, 28 have not been described in the current published literature as autoantigens associated with autoimmune disease and are potentially novel biomarkers **(Supplementary Table 5**). These results show multiple proteins from the XIST RNPs are novel autoantigens in patients with DM, SSc, and SLE.

Finally, we turned to our mouse model to probe the drivers of autoantibodies to Xist RNP. Using our Xist RNP array, we analyzed sera from the tgXist mice of the SJL/J background to examine the effect of Xist expression and pristane induced lupus. We compared sera of each animal longitudinally before and after 16 weeks of Xist induction or pristane treatment, allowing us to infer a causal relationship between perturbation and autoantibodies to Xist RNP. Moreover, we compared the autoantibodies in tgXist mice to autoantibodies to the same proteins in human patients with SLE, which grounds the mouse results with human disease relevance (revised **Fig. 7A**). Pristane treatment of WT female mice induced autoantibodies to 41 Xist RNP proteins, 26 of which are shared with SLE patients, including the high stringency XIST-binding proteins HNRNPK, HNRNPUL2, SF1 and WAC (**Figure 7B**). Importantly, induction of tgXist expression in male mice in the absence of pristane (tgXist M) was sufficient to induce autoantibodies to 44 Xist RNP proteins, of which 27 were shared with SLE patients (*p*= 0.003 for overlap, **Fig. 7B**). This result indicates that Xist expression and RNP formation in male tissues suffice to initiate autoantibodies with homeostatic turnover. Finally, 30 of 45 enriched protein antigens from pristane-treated tgXist-expressing male mice overlapped with SLE patient enrichment lists (*p*= 0.001 for overlap, **Figure 7B**). These results suggest that Xist can promote autoantibodies to Xist RNP which precedes disease onset, and that many of the autoantibodies are likely biomarkers of the disease but may not be causal in the absence of tissue damage.

**Figure 7.**
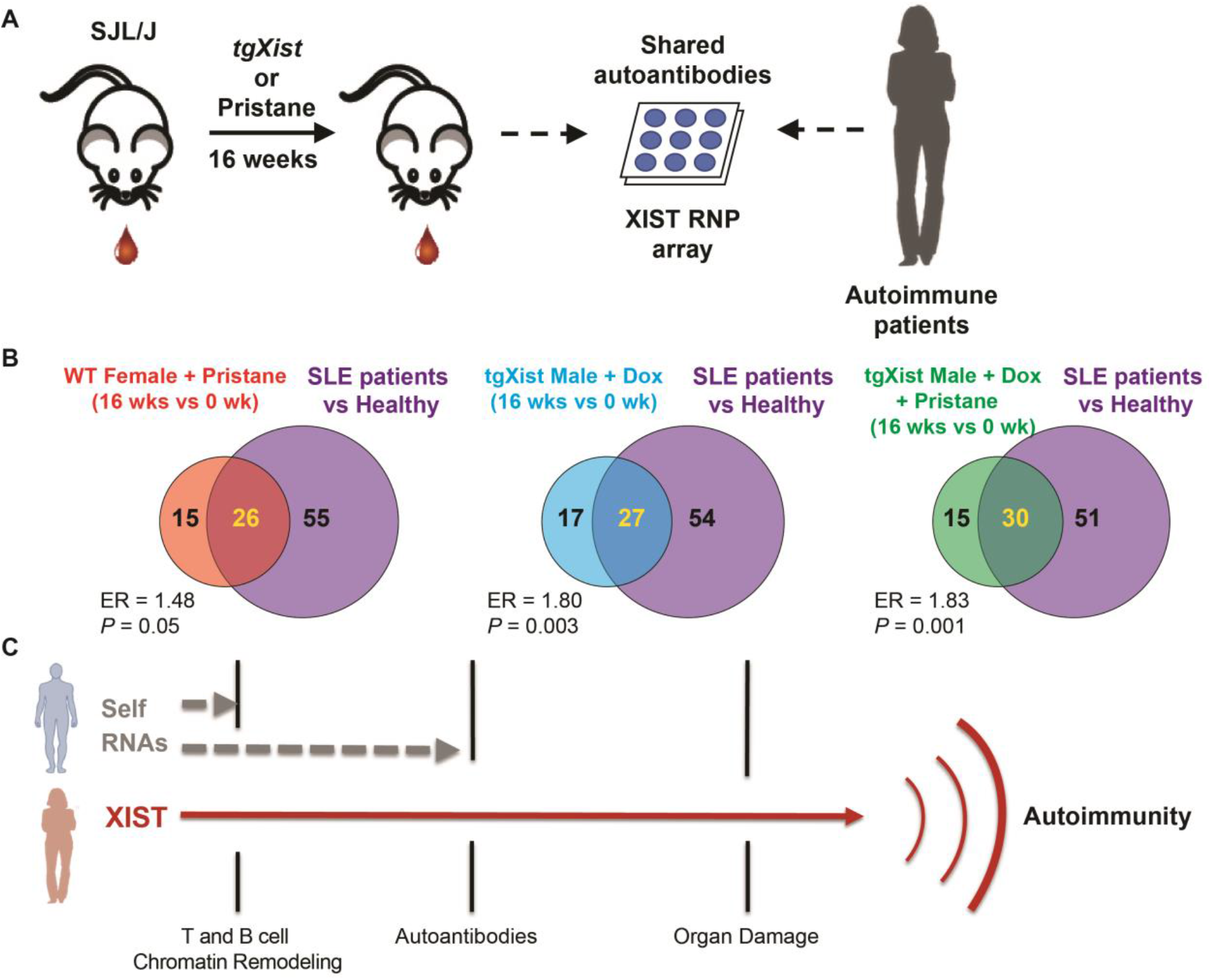
Model of XIST RNP in autoimmune progression. (A) Schematic using the Xist antigen array to identify autoantibodies shared in SLE patients and *tgXist/Xist*-expressing SJL/J mice with/without pristane-induced SLE. (B) Metrics of autoantibodies from the *Xist* antigen array enriched in SLE patients compared to the general population and tgXist mice 16 weeks after pristane injection and/or induction of *tgXist* compared to treatment baseline. ER= enrichment ratio, significance calculated using the Fisher’s Exact Test (Wt Female + pristane n=13, tgXist Male + Dox n=11, tgXist Male + Pristane + Dox n=9). (C) Autoreactivity to *Xist* RNPs first causes changes in the chromatin landscape impacting genomic accessibility and changes in lymphocyte gene expression programs prefacing the development of autoantibodies and cascade to prolonged autoreactive activity damaging organs in the final stages of autoimmunity.

## DISCUSSION

### Xist lncRNA as a polymeric antigen scaffold in female-biased autoimmunity

Our study nominates *Xist* ribonucleoprotein complexes as antigenic triggers underlying the greater prevalence of autoimmune diseases in females. Although it is a well-documented fact that females are more prone to autoimmune diseases than males, previous studies primarily examined differences in gene contribution and hormonal background. While recent studies of *Xist* address altered X-inactivation and the subsequent impact of XCI escape of X-linked genes^14,59,60^, this study is the first to investigate the immunogenicity of *Xist* RNP complex itself. We have shown that expression of Xist RNPs in male mice is sufficient to increase disease severity and change the expression and epigenomic profiles of both the B-cell and T-cell effectors of SLE pathogenesis.

Physicians and scientists have long noted that many autoantibodies target large nucleic acid protein complexes, such as chromatin or RNP, in human autoimmune diseases. This feature was exploited by molecular biologists to use patient sera to identify components of the centromere (recognized by autoantibodies in CREST syndrome) or spliceosome (SLE, DM). Immunologists have explained this phenomenon with the idea that large nucleic acid-protein complexes are polymeric, and if exposed in the extracellular space, can cluster and activate immunoreceptors. We propose that the XIST RNP is one such dominant antigenic array that is unique to females. Every cell in a woman’s body has XIST, which is a long polymer (19 kb) and coats the entire inactive X chromosome in the condensed Barr body (an even larger polymer). When a female cell dies due to tissue injury, XIST RNPs will invariably be exposed to the immune system. Our data further suggests a model where XIST contributes to several steps in the progression to autoimmune disease (**Figure 7C**). In a genetically autoimmune resistant background, low level of XIST, even in the presence of tissue injury, leads to only autoantibody production but no frank organ pathology. These incipient changes are associated with changes in T cell subsets and chromatin state changes toward more effector T cells. These epigenetic changes in accessibility are then subsequently reflected in the gene expression programs upregulating autoreactivity and downregulating immune modulation. Finally, in the context of a permissive genetic background and repeated tissue injury, the presence of XIST RNP exacerbates full blown end organ pathology and activation of multiple immune cell types. Longitudinal studies of sera reactivity and autoimmune disease in humans are consistent with this model ^61^.

### Opportunities for disease diagnosis and therapy

There are more than 100 known autoimmune diseases that, in aggregate, combine to afflict as many 50 million Americans and comprise one of the top ten leading causes of death for women under the age of 65^62^. Worryingly, cases are increasing yearly on a global scale and recent serologic studies revealed a steep rise of increasing ANA reactivity^63,64^. Understanding the risk factors and drivers of autoimmunity has become even more critical in the race to develop effective therapies and sensitive diagnostics specific to each autoimmune disease. However, the high heterogeneity within autoimmune diseases and overlapping traits across diseases have limited our ability to tailor effective therapies and sensitive diagnostics specific to each autoimmune disease^65^. Our discovery of seropositivity towards multiple XIST-associating proteins in autoimmune patients introduces a novel antigen set with great clinical potential for enhancing diagnostics and disease monitoring, as autoantigens are often detected prior to or early in disease onset^61,66^. In addition to serving as effective biomarkers for disease, studies in SSc have also demonstrated the effectiveness of autoantigen analysis in patient stratification and identifying pathogenic pathways^67,68^. Profiling the XIST RNP in primary cells, both healthy and diseased, will also be essential in advancing our understanding of the aberrant autoreactivity towards proteins within the complex and identify more potential autoantigens.

Currently, there are few targeted therapies for autoimmune diseases available. The most common therapies involve B-cell depletion, but are not always effective. There remains a need for more specific pathogenic leukocyte targets. We identified atypical B cells as a population of immune cells that accumulate as a consequence of Xist RNP expression. Atypical B cells (also known as age-associated B cells) are a unique population of B cells that expand with increased TLR7 signaling^69,70^ and in female-biased autoimmunity^71^. Notably, atypical B cells accumulate in aged female mice but not in age-matched male mice^69^, and atypical B cells are enriched in human or mouse B cells that escape XCI and re-express *TLR7*^14,72^. Thus, atypical B cells appears as the immunological nexus of two potential consequences of mammalian dosage compensation--autoreactivity to Xist RNP and escape from XCI. Identifying novel antigens and their associated pathogenic pathways hold great therapeutic potential. Future studies should address whether and how atypical B cells or other cell types evoked by Xist RNP contribute to autoimmunity.

### Limitations of the study

In this study, we deployed a transgenic *Xist* allele that lacked the A-repeat. Therefore, we could not assess the potential contribution of proteins that bind to the A-repeat, such as Spen, in autoimmunity. Our transgenic model expressed *Xist* ubiquitously in male animals, and the role of individual tissues or cell types most responsible the observed phenotypes and the required time window of Xist RNP exposure should be dissected in future studies. Additionally, a known limitation of Tet-regulated expression cassettes is that they can be silenced over time *in vivo*^73–76^. Thus, there may be variability or decline of dox-induced *tgXist* expression in our transgenic model over the experiment. In a related aspect, this work does not address whether fluctuations in Xist level in female individuals may impact autoimmunity.

## ACKNOWLEDGEMENTS

We thank members of the Chang and Utz labs for discussion and Adrianne Woods and Gwendolyn Leatherman for assistance with serum compilation from Johns Hopkins. We thank Greg S. Nelson for assistance with mouse blood collection, the Stanford Veterinary Service Center and the Stanford Breeding Colony Management Services for assistance with animal husbandry and welfare, the Stanford Comparative Medicine Services for histology preparation, the Stanford Transgenic, Knockout and Tumor Model Research Center for tissue sample preparation, the Stanford Functional Genomics Facility for bulk sequencing assistance, and the Stanford Shared FACS Facility for assistance in cell sorting. This work was supported by Scleroderma Research Foundation (H.Y.C.), NIAMS T32 AR007422 and NIAMS K99/R00 (D.R.D.), and NIAMS T32 AR050942 (B.T.A.). The Hopkins Lupus Cohort is supported by a grant from the National Institute of Arthritis and Musculoskeletal Diseases under award R01-AR069572. H.Y.C. is an Investigator of the Howard Hughes Medical Institute.

## AUTHOR CONTRIBUTIONS

Conceptualization: D.R.D., P.J.U., and H.Y.C.; Methodology and Investigation: D.R.D, K.M.C., R.L., D.C., K.K., C.H., R.S., S.C., A.F., D.W.G., A.A.S., M.P, F.M.W., L.S.C., and D.F.F., and A.W.; Data Analysis: D.R.D., Y.Z., J.A.B., S.S., Y.Z., K.M.C., B.Y., B.A., C.H., R.S., A.F., B.A., and E.K.L.; Writing: D.R.D., K.M.C., L.S.C., D.F.F., P.J.U., and H.Y.C.; Funding Acquisition: H.Y.C.; Resources: H.Y.C.; Supervision: H.Y.C.

## MATERIALS AND METHODS

### Bibliomic Analysis

In total, 118 XIST-associated proteins published as high confidence XIST RNP proteins^14^ were reviewed for known autoantigen status. The bibliomic analysis was performed by primarily mining a human autoantigen database of peer-reviewed articles^77^ and additional manual aggregation. Each table entry, represented as ^75^ ^Excerpt^, was curated to remove false positives resulting from autoantigen and autoantibody studies of diseases such as cancers, paraneoplastic syndromes and non-specific/inconclusive results. The mined data was further organized to include the most relevant excerpt from each article associated with a single PubMed ID and thereby avoiding duplicate entries. Additionally, missing and ambiguous “Disease” terms were resolved with domain-based imputation^78^.

From the 118 high confidence XIST-RNP proteins, bibliomic analysis collated data on 30 XIST proteins, across 57 disease terms and 451 unique PubMed articles (**Supplementary Table 1, Supplementary Figure 1**).

#### XIST autoantigen array additional proteins

Among the reactive proteins found in patient serum from the XIST antigen array, bibliomic analysis identified 2 additional XIST associating proteins across 5 diseases and 8 unique PubMed articles (**Supplementary Table 1**).

### Mice

#### Strains

Wild-type C57BL/6J (000664) and SJL/J (000686) mice were purchased from Jackson Laboratories. Inducible **Δ**RepA-Xist transgenic mice: A transgene of tetO linked to *Xist* cDNA deleted for the A repeat (SacII-XhoI is deleted, appx. 500nt) was targeted to the *Col1a1* locus on chromosome 11 in A9 129/Bl6 hybrid ES cells. To insert the **Δ**RepA-Xist transgene, a homing cassette was first inserted into the 3’-region of *Col1a1* using homologous recombination. This homing cassette consisted of a hygromycine (pPGK-hygro-PA) resistance marker, a loxP recombinase site, and a truncated neomycine resistance gene (3’neo-PA). Subsequently, the pPGK-loxP-**Δ**RepA-Xist cDNA was inserted by Cre mediated recombination followed by G418 selection^18,79^. Transgenic mice were generated by injection of the modified ES cells into Bl6 8-cell embryos^80^. Resulting transgenic mice were crossed to a mouse line carrying the R26/N-nlsrtTA doxycycline regulated transactivator^80,81^. Mice carrying the tet-O-ΔRepA-Xist and rtTA constructs were backcrossed into the C57BL/6J or SJL/J mice for multiple generations. Tail tips were collected from 5-7 female mice from each generation for speed congenics selection using Charles Rivers’ MAX-BAX 384 SNP panel. The two females with the highest match in the desired background from each generation were selected as breeders. The tgXist-C57BL6/J mice used in this study are > 80% in the C57BL/6J background. The tgXist-SJL/J mice are > 99.99% in the SJL/J strain background. Mice were genotyped for both the Neomycin resistant cassette and from the CMV promoter into the truncated Xist transgene. Studies in the SJL/J background used both mice heterozygous for tgXist and wild-type littermates while studies involving SJL/J mice while wild-type control C57BL/6J mice were obtained from Jackson Laboratories. Genotyping primers are listed in **Supplementary Table 8**.

#### Pristane-induction of SLE

Wild-type and tg-Xist mice were injected with a one-time injection of 0.5 mL of pristane (Sigma-Aldrich, P9622-10X1ML) at 8-10 weeks old (SJL/J strain) or 12-14 weeks old (C57BL/6J studies). Control animals were injected with PBS 1x (Thermo Fisher Scientific, 10010049).

#### Induction of transgene (tgXist)

Simultaneous to the injection date, doxycycline (Sigma-Aldrich, S9891-5G) was continuously administered at 0.2 g/mL in drinking water to select mice until the terminal timepoint.

For the *tgXist* expression validation studies, *tgXist^+/-^* and wild-type mice in the C57BL/6J strain were administered doxycycline for 2 weeks and tissues were harvested for qRT-PCR and FISH analysis.

#### Xist qRT-PCR

Mechanically dissociated thymus, spleen, kidney, and liver cells harvested from *tgXist^+/-^* and wild-type mice in the C57BL/6J were strained through a cell filter, pelleted, and frozen in RLT Buffer with 0.1% BME. RNA was extracted using the RNeasy Mini kit (Qiagen, 74106), genomic DNA was removed using amplification grade DNAse I (Thermo Fisher Scientific, 18068015). RNA concentration was quantified on a Nanodrop. cDNA was made from 1 ug of RNA/sample using the High-Capacity cDNA Reverse Transcription Kit (Applied Biosystems, 4368814). Each Ct value was measured using Lightcycler 480 (Roche) and each mean dCt was averaged from triplicate qRT-PCR reactions. Relative Xist RNA levels was calculated by ddCt method compared to GAPDH controls. Statistical significance was calculated using the Student’s t-test. Primer sequences are listed in the **Supplementary Table 8**.

#### Xist FISH

Mice organs frozen in O.C.T. compound were sectioned on a crysostat to 15 µm onto microscope slides (Fisher Scientific, 12-550-15). Sections were washed once with PBS 1x, fixed in 4% paraformaldehyde for 10 min at room temperature, washed twice with PBS 1x for 2-5 min each, then permeabilized on ice with ice cold 0.5% TritonX-100 in PBS for 10 min. Slides were then rinsed once with PBS and dehydrated sequentially in 70%, 90% and 100% Ethanol (Gold Shield, 412804) for 5 minutes each. Sectioned slides were then allowed to air dry before immersion in freshly made Stellaris® RNA FISH Wash Buffer A (SMF-WA1-60) for 2-5 minutes at room temperature. Sections were then hybridized overnight for 16-18 hours in the dark at 42C in 250 nM of Mouse Xist Stellaris® FISH Probes with Quasar® 570 Dye (SMF-3011-1) in Stellaris® RNA FISH Hybridization Buffer (SMF-HB1-10). The next day, slides were incubated twice in Wash Buffer A for 30 minutes at 37C followed by immersion in Stellaris® RNA FISH Wash Buffer B for 2-5 minutes at 27C, mounted in VECTASHIELD Mounting Medium with DAPI (Vector Laboratories, H-1200) and sealed with cover glass (VWR, 48366-227) and nail polish clear coat. Sectioned organs were imaged on the Zeiss Observer Z.1 using the 63x oil objective, X-Cite Series120 laser, and AxioVision Rel. 4.8 software.

#### Mouse sera collection

Mice were sedated with isofluorane and retro-orbitally bled immediately prior to the injection and viably bled every 4 weeks until the terminal date. On the terminal date, blood was collected through cardiac puncture. Blood was allowed to clot for 2 hours at room temperature, then spun for 15 minutes at 1500xg at room temperature. Serum was immediately collected, flash-frozen on dry ice and stored at -80C.

#### ANAb ELISA

Serum ANAb levels were measured using the Mouse Anti-Nuclear Antibody Kit (MyBioSource.com, MBS731183) and assessed on an ELISA plate reader according to manufacturer’s protocol. Titration curves and values were calculated using a Four Parameter Logistic Curve.

#### Tissue collection and preparation of pristane-induced SLE studies

Mice were euthanized by CO_2_ asphyxiation and cardiac exsanguination at the terminal timepoints of 16- or 52-weeks post-injection. Terminal cardiac blood was aliquoted into EDTA tubes (400 ml per mouse) and Eppendorf tubes (for serum collection). Total body weights were obtained, and the following organs were weighed individually: thymus, heart, liver, spleen, testes. Gross necropsies were performed on all mice (**Supplementary Table 9**). The caudate and papillary liver lobes, left kidney, one half of the thymus and one half of the spleen were dissected and placed on ice in HypoThermosol FRS (BioLifeSolutions, 101102) until all dissections concluded and were either mechanically dissociated using clean scalpels and syringes or frozen in O.C.T. Compound (Tissue-Tek, 4583) at -80C on the same day. The remaining tissues were immersion-fixed in 10% neutral buffered formalin for 72 hours for downstream histology analysis. Dissociated cells were passed through cell filters (Sysmex, 04-004-2327), pelleted at 300xg, incubated for 1 minute at room temperature in eBioscience™ 1X RBC Lysis Buffer (00-4333-57) to remove red blood cells, and quenched with 10x the volume of PBS 1X. For viable stocks, dissociated cells were frozen in BAMBANKER (Wako, #203-14681) and stored in liquid nitrogen. For RNA extraction, cells were frozen in either TRI Reagent® (Molecular Research Center, TR118) or Buffer RLT Plus (Qiagen, 1053393) and stored at -80C.

#### CD4+ isolation

CD4+ cells were isolated from freshly dissociated mouse spleen using the EasySep Mouse CD4+ T cell Isolation Kit (STEMCELL Technologies, 19852). CD4+ cells were then immediately used for bulk ATAC-seq, frozen in RLT Buffer Plus, or viably frozen in BAMBANKER. A small aliquot of cells were stained in PBS 1x with the viability marker 7-AAD (BD Biosciences, 559925) and T-cell marker CD4 FITC (clone GK1.5, BioLegend, 100406) to assess purity and viability on a FACS analyzer.

#### CD45+ isolation

Viable cells frozen in BAMBANKER were thawed at 37C in a bead bath, resuspended in RPMI, and spun at 300xg to pellet and remove the buffer. Cells were stained with the pan-hematopoietic marker CD45 APC (clone 30-F11, BioLegend, 103112) and 7-AAD for viability. Viable CD45+ cells were sorted on the BD FACSAria II sorter using a 70 uM nozzle into chilled FBS in the Stanford Shared FACS Facility. Sorted cells were then immediately used to prepare sequencing libraries.

#### Histopathology

Formalin-fixed tissues were processed routinely, embedded in paraffin, sectioned at 5 µm, and stained with hematoxylin and eosin. All tissues were evaluated blindly by a board-certified veterinary pathologist (KMC). An ordinal histopathologic grading scale (score 0-4) was designed to evaluate glomerulonephritis, hepatic lipogranulomas, pulmonary lymphohistiocytic alveolitis, and splenic extramedullary hematopoiesis. A binary histopathologic grading scale (score 0 or 1) was used to evaluate for intraabdominal lipogranuloma formation and ectopic lymphoid tissue, pulmonary hemorrhage, hemosiderosis, and/or vascular thrombosis, splenic lymphoid hyperplasia and plasmacytosis, and lymph node hyperplasia and medullary plasmacytosis. A total composite score was derived for each mouse. The Fisher’s Exact Test was used to calculate significance between treatment groups. Due to facility and equipment difficulties during the COVID-19 pandemic, some mice are missing scores for kidney or spleen. Only mice with all organs correctly processed and assessed were used to calculate the composite score and included in the final analysis (shown in **Figure 3**). Complete scores for all SJL/J mice used in the pristane-induced lupus study can be found in the **Supplementary Table 9**.

#### Statistical Analysis of histopathology scores

The Fisher’s Exact Test was used to calculate significance between treatment groups in the histopathology studies. Because the lowest positive female control total body disease score was 10, we set 10 as the cutoff total body disease score to distinguish between low disease and high disease scores.

### Sequencing Library Preparation

#### Bulk sequencing

ATAC-seq libraries of freshly isolated CD4+ cells were prepared using the Omni-ATAC protocol^82^. RNA was extracted using the RNeasy Plus Mini Kit (Qiagen, 74136). TruSeq® Stranded mRNA Library Prep (Illumina, 20020594) was used to generate polyA-selected RNA-sequencing libraries, cleanup performed on magnets with AMPure XP beads (Beckman Coulter, A63880). The bulk ATAC-seq and RNA-seq libraries were sequenced with paired-end 75 bp read lengths on in the Stanford Functional Genomics Facility an Illumina HiSeq 4000 that was purchased with funds from NIH under award number S10OD018220.

#### Single cell sequencing

The Chromium Single Cell Multiome ATAC + Gene Expression (10x Genomics) was used to prepare libraries for CD45+ sorted cells with a target of 10,000 cells/sample. Libraries were sent to Novogene for Bioanalyzer trace quality control check and sequencing. Libraries were sequenced on the NovaSeq 6000 at a depth of 20,000 paired reads per cell for gene expression libraries and 25,000 paired reads per cell for ATAC libraries.

### Sequencing Library Analysis

#### Bulk sequencing

The bulk RNA-seq data was aligned to mm10 using STAR. The gene expression read counts was generated using RSEM. The adaptor of paired-end ATAC-seq data were first trimmed by an in-house software, and then aligned to hg38 genome using bowtie2. The mitochondrial reads and reads with low alignment score (<10) were removed. The aligned sam files were converted to bam files and sorted by Samtools. Picard was used to remove duplicate reads and MACS2 was used to call peaks. Each ATAC-seq peak was annotated by its nearby genes using GREAT under the basal plus extension default setting.

The raw bulk RNA-seq and ATAC-seq read counts were then normalized and analyzed using DESeq2. The differentially expressed genes and ATAC-seq peaks were identified using the negative binomial models. Benjamini hochberg procedure was used to adjust for multiple hypothesis testing. Peaks with FDR < 0.2 and absolute fold change larger than 1.5 were selected as significant. Bioinformatics tool g:Profiler was used for pathway enrichment analysis. CIBERSORT was used to estimate the abundance of immune cells based on normalized RNA-seq data.

#### Single cell multiomics sequencing

The single-cell paired RNA and ATAC-seq reads were aligned to the mm10 reference genome using cellranger-arc count (10x Genomics, version 2.0.1).

Gene expression data was filtered to include only barcodes that had nFeature > 500 and pct.mt < 12. Two female samples (F100 and F113) were excluded because, even after stringent filtering, cells from these samples had lower quality metrics than other samples. R v4.1.3 Seurat v4.1.1 and ggplot2 v3.3.6 were used for downstream analysis and visualization. After filtering, clustering and dimensionality reduction was performed using the top 10 principal components (dims=1:10) and a clustering resolution of 0.25. Differential gene expression analysis was performed using Seurat function "FindMarkers" with default settings.

Single cell ATAC data was processed with ArchR v1.0.1. ATAC-seq data was subsetted using the subsetArchRProject function to include only cells with matched gene expression cell IDs and clustered at resolution=0.7 with default settings. Gene imputation features of defining cell type markers were visualized from the GeneScoreMatrix using default settings. Cellular subsets were subsequently subsetted and analyzed with getMarkerFeatures from the PeakMatrix using default settings and maxcells=5000 and k=1000.

Quality Control metrics for the Multiome libraries can be found in **Supplementary Table 10**.

#### Calculation of Chr11 accessibility and gene expression

The aforementioned Chromium Single Cell Multiome ATAC + Gene Expression (10x Genomics) libraries were used in the analysis of Chr11 between the three treatment groups (Wt F+ Pristane, tgXist M + Pristane + Dox, and Wt M + Pristane + Dox).

#### Chr11 track analysis

The bigwig files for each mouse treatment group were generated using the getGroupBW function in the archR package. Boxplots were visualized in the rtracklayer package in R using bigwig files containing normalized ATAC-seq counts. Genome position-wide scATAC track visualization was obtained using the plotBrowserTrack function in the archR package. The FindMarkers function in the Seurat package was used to identify the differential expressed genes between the pristane+dox treated groups: Wt M versus and tgXist M. A hypergeometric test was then applied to examine whether the genes on chr11 are more likely to be down regulated compared with the genome wide scale.

### Clinical Cohorts

All human patient samples were de-identified in this study.

#### General Population

Donated whole blood from a total of 9 male and 8 female donors were obtained from the Stanford Blood Bank following IRB-approved protocols with confirmed assent for use in research. Serum was collected as described above in the mouse serum section and frozen in -80°C for long-term storage. During data analysis, 1 of the 9 male samples was filtered out for high IgG background.

#### Dermatomyositis (Stanford)

Serum from 1 male and 3 female dermatomyositis patients were used in this study. All samples were collected from patients seen at the Stanford outpatient clinics under an IRB-approved protocol, and all patients provided informed consent to participate. The dermatomyositis cohort has been described previously^83^. All patients met probable or definite DM by 2017 ACR/EULAR IIM Classification Criteria^84^. All sera used in the study were also known to contain antibodies against TIF1-γ which was assayed as previously described^85^.

#### Systemic Sclerosis (Scleroderma, Stanford)

Serum from 24 patients (2 male and 22 female) with systemic sclerosis (SSc) were included in this study from the Stanford cohort. All samples were collected from patients seen at the Stanford outpatient clinics under an IRB-approved protocol, and all patients provided informed consent to participate. All patients fulfilled 2013 ACR/EULAR classification criteria for SSc. Twelve patients had diffuse and 12 had limited cutaneous SSc. Seven patients had the Scl-70 antibody, 5 had the anti-centromere antibody, 2 patients had RNA polymerase III, 2 had nucleolar ANA, 2 had PM/Scl, and 1 had U1RNP, with the remainder having positive ANA but no known SSc-specific autoantibody.

#### Systemic Sclerosis (Scleroderma, Johns Hopkins)

Serum from 9 males and 31 females were shipped on dry ice to Stanford. Patients were enrolled in the IRB-approved Johns Hopkins Scleroderma Center Research Registry. Scleroderma participants in the registry meet at least one of the following criteria for systemic sclerosis: 1) 2013 ACR/EULAR classification criteria for scleroderma, 2) 1980 ACR classification criteria, 3) having at least 3 of 5 features of the CREST syndrome (calcinosis, Raynaud’s phenomenon, esophageal dysmotility, sclerodactyly, telangiectasia), or 4) having definite Raynaud’s phenomenon, abnormal nailfold capillaries and a scleroderma-specific autoantibody.

#### Systemic Lupus Erythematosus (Johns Hopkins)

Patients provided written informed consent to participate in the Hopkins Lupus Cohort (IRB study number NA_00039294). Blood was drawn at the time of a clinical blood draw, serum collected and kept in a -80 freezer for long term storage. For this study, 40 serum samples (3 males and 37 females) were randomly selected from patients who were ever: 1) ANA positive with a titer of at least 1:320; and 2) positive to the dsDNA autoantigen. Patients enrolled in the Hopkins Lupus Cohort met either the revised American College of Rheumatology (ACR)^86^ or Systemic Lupus International Collaborating Clinics (SLICC)^87^ criteria for SLE. Samples were shipped on dry ice to Stanford and stored at –80°C.

### SLE autoantigen Panel Array

#### Assay

Mouse serum was assessed using the SLE autoantigen panel and array as previously described^36^. Serum from the baseline timepoint of 0 weeks (injection start) and terminal timepoints (16 weeks for SJL/J and 52 weeks for C57BL/6J strains) were run in the same assay (sample-matched). Serum was stored long-term at –80°C and prepared as previously described^36^. Raw MFI values and mouse serum sample information can be found in **Supplementary Table 2**.

#### Analysis

Raw MFI scores were first normalized by subtracting the baseline value of bare bead. The difference between terminal and baseline timepoints was then calculated using the normalized values. Values were plotted in GraphPad PRISM and the Student’s t-test was used to test for statistical significance.

### XIST Autoantigen Array

#### Autoantigen list

Available recombinant proteins were chosen from a set of XIST-associating proteins of interest from published XIST ChIRP datasets^12,14^ at a stringency of log2(EXP/RNAse) > 1 and peptide EXP Sum ≥ 10. Due to the limited availability of recombinant proteins, this criteria is less stringent than that used for the bibliomics analysis in order to include a larger set of XIST-associating proteins of interest. Autoantigens clinically used to screen for DM^53^, SLE or SSc^88,89^ were included as positive controls. 16 exploratory proteins were included from unpublished XIST ChIRP-MS lists. Each protein was represented by one or more protein fragments (20-151 amino acids long) produced within the Human Protein Atlas project (https://www.proteinatlas.org/)^90,91^. The full list of autoantigens and their categorization can be found in **Supplementary Table 3**.

#### Sample preparation

25 uL of serum per sample were aliquoted in a pre-determined randomized order onto 96-well plates (Thermo Fisher Scientific, AB800150) with mouse and human samples on separate plates. Thermal sealed plates were shipped on dry ice to SciLifeLab in Sweden and stored at – 80C upon arrival.

#### Suspension Bead Array Assay (SciLifeLab)

The antigens were immobilized on color coded magnetic beads (MagPlex, Luminex Corp., Austin, TX) and the assay was run as previously described, with minor alterations^92^. The samples were diluted directly before running the assay, prior to adding the secondary antibody the beads incubated for 10 minutes in 0.2% paraformaldehyde to fixate any bound antibodies, and the secondary antibody used was R-Phycoerythrin labelled Goat anti-Human IgG Fc (eBioscience^TM^; 12-4998-82, Invitrogen).

#### Data handling and analysis

All data processing from the suspension bead array was performed using R version 4.1.1. Antigen specific background was adjusted for by centering and scaling the 10^th^ percentile of the mean fluorescence intensity (MFI) value for each antigen to a common value (10^th^ percentile of the whole dataset). The antigen percentile adjusted MFI values were transformed per sample into the number of “median absolute deviations” (MADs) around the sample, represented as: MADs_ag,sample_ = (MFIadj_ag,sample_ − median_sample_) / MAD_sample._

For the general population and patient disease comparison analyses, the difference in reactivity was calculated as the difference between the mean of the MAD scores of the comparison groups and significance was calculated using the Student’s T-Test. To remove far-lying outliers, MAD scores were first “trimmed” to quantile (0.1, 0.9). Since each protein often contained multiple protein fragments in the assay, the list was next filtered to remove duplicate counts and generate a unique list reactive proteins. The reactive protein lists were then compared between patient disease groups (**Figure 6D,E, Supplementary Table 5**).

#### Selection of enriched proteins

To find shared reactive proteins in two heterogeneous datasets (pristane-induced SLE mouse and autoimmune disease patients), the difference in MAD scores was calculated in patients as the difference between SLE patient reactivity and the general population (healthy control) and in mice as the difference between 16 weeks and 0 weeks (treatment start baseline) for each group. Protein fragments were considered reactive in patients if MADs_diff_ > 0 in the quantile “trimmed” set and MADs_diff_ > 2.5 in the raw datasets. Protein fragments were considered reactive in mice if MADs_diff_ > 2.5 between 16 and 0 weeks. Since each protein often contained multiple protein fragments in the assay, the list was next filtered to remove duplicate counts and generate a unique list reactive proteins. The enriched protein lists were then compared between the mouse groups and the SLE patients (**Figure 7B**, **Supplementary Table 7**).

