## Supplementary Figures 1, 2, 3, 4, 5, 6 and Supplementary Table captions for "Xist ribonucleoproteins promote female sex-biased autoimmunity"

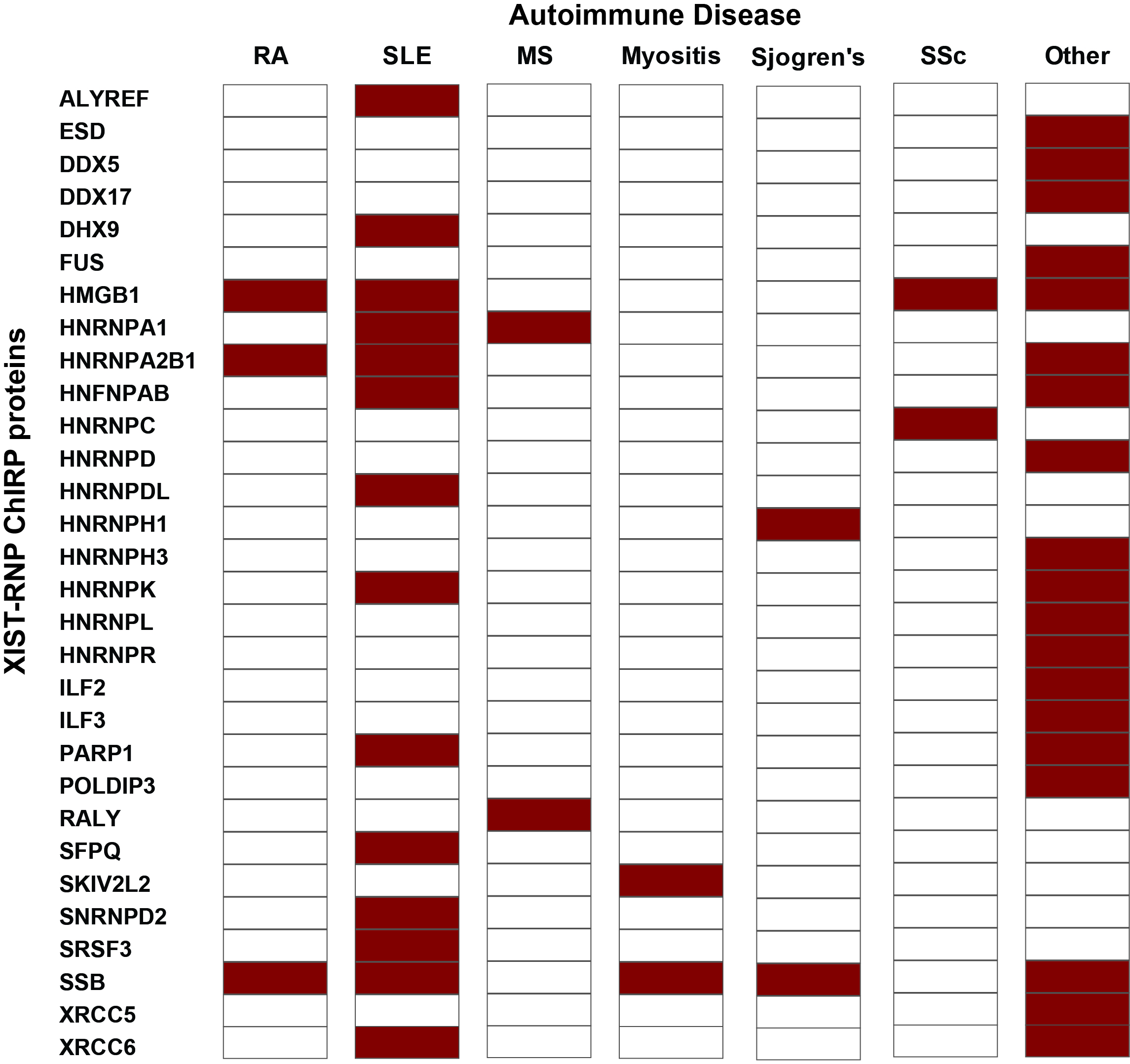

**Supplementary Figure 1: Established autoantigenic associations of XIST RNPs.** Summary of XIST RNP complex proteins with known association as autoantigens in autoimmune disease grouped by disease (Red=positive hit). Full bibliomics information available in **Supplementary Table 1**.

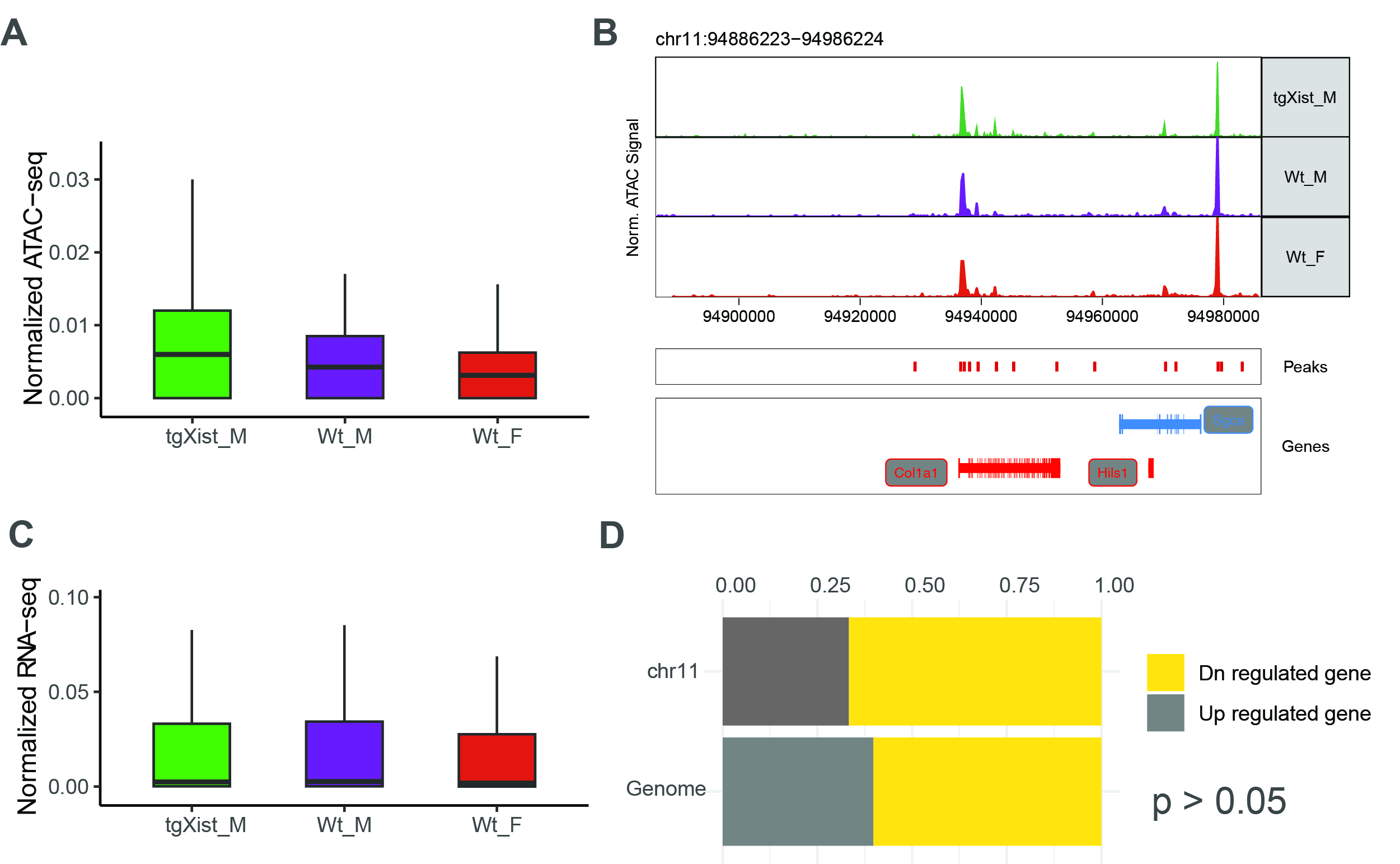

**Supplementary Figure 2: Chromosome-wide comparison of tgXist transgene insertion in Col1A1 safe harbor locus on Chr11.** (A) Chromatin accessibility at Chr11 assessed using single cell ATAC-seq data (B) ATAC-seq accessibility tracks in the Col1A1 insertion locus (C) Gene expression changes at Chr11 compared to the whole genome (D) Fraction of up- and down-regulated genes on Chr11 and genome-wide comparing tgXist males and Wt male mice. All plot values obtained from single cell multiomic data normalized to sequencing depth from pristane-treated SJL/J mice: number of tgXist M+ Dox + Pristane= 8, Wt M+Dox+Pristane= 3, Wt F + Pristane= 2. Significance was calculated using Fisher’s Exact test.

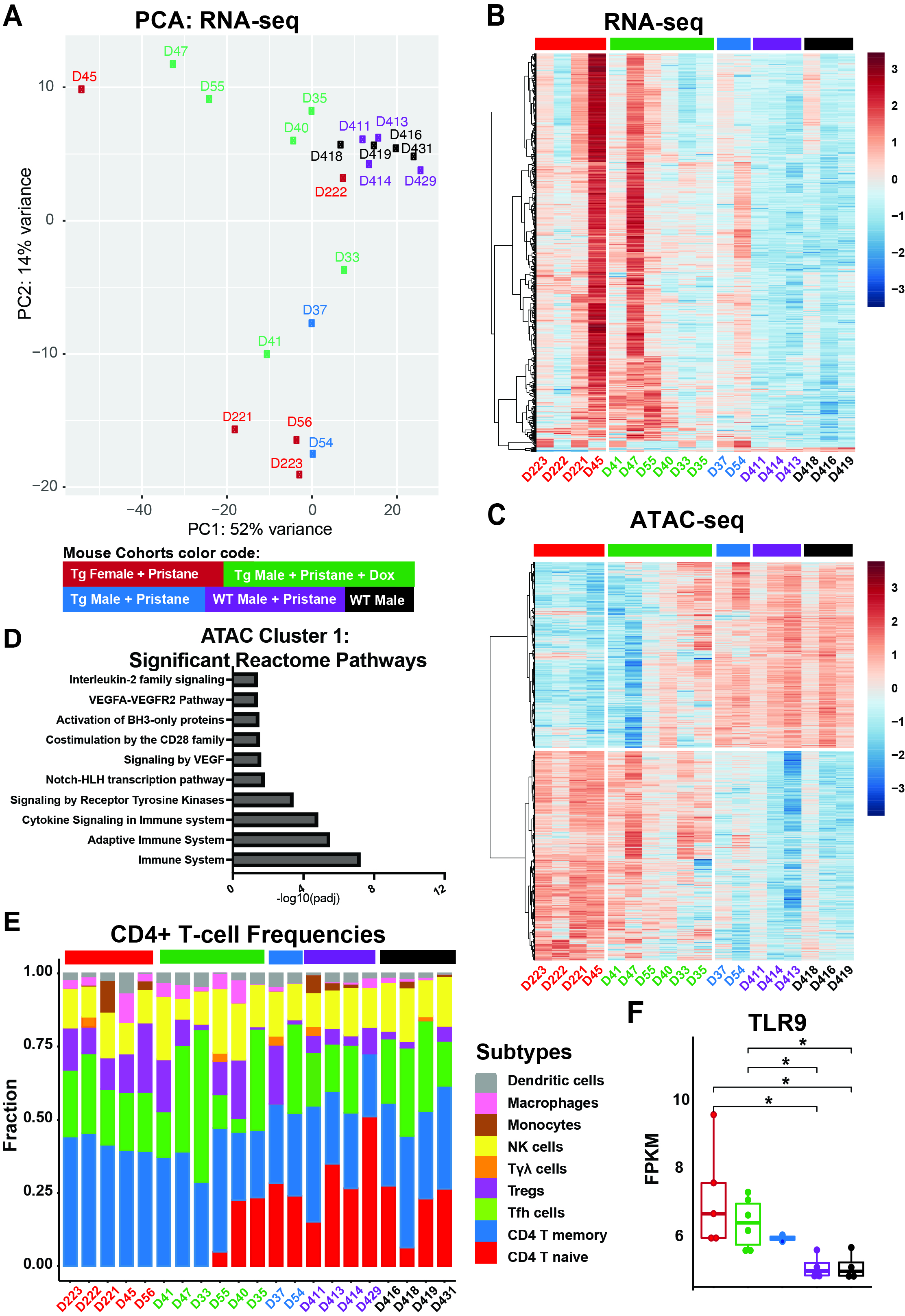

**Supplementary Figure 3: CD4+ Splenic T-cells ATAC- and RNA-sequencing from pristane-induced SLE mice of the C57BL/6J strain**: (A) PCA plots of RNA-seq libraries of all five mouse cohorts. (B) RNA-seq heatmap of differentially expressed genes and (C) ATAC-seq of differential peaks from complete group of mice used in the bulk sequencing analysis. (D) Top 15 differential reactomes associated with ATAC-seq cluster 1 genomic regions. (E) CIBERSORT deconvolution of CD4+ subtypes of all mice used in the bulk sequencing analysis. (F) Comparison of Fragments Per-Kilobase per-Million mapped fragments (FPKM) in the TLR9 gene region. Significance calculated from Student’s T-test. Number of mice used: WtF+Pristane=5, tgXist M+Pristane+Dox=6, tgXist M+Pristane=2, Wt M+Pristane=4, Wt M mock treatment=4.

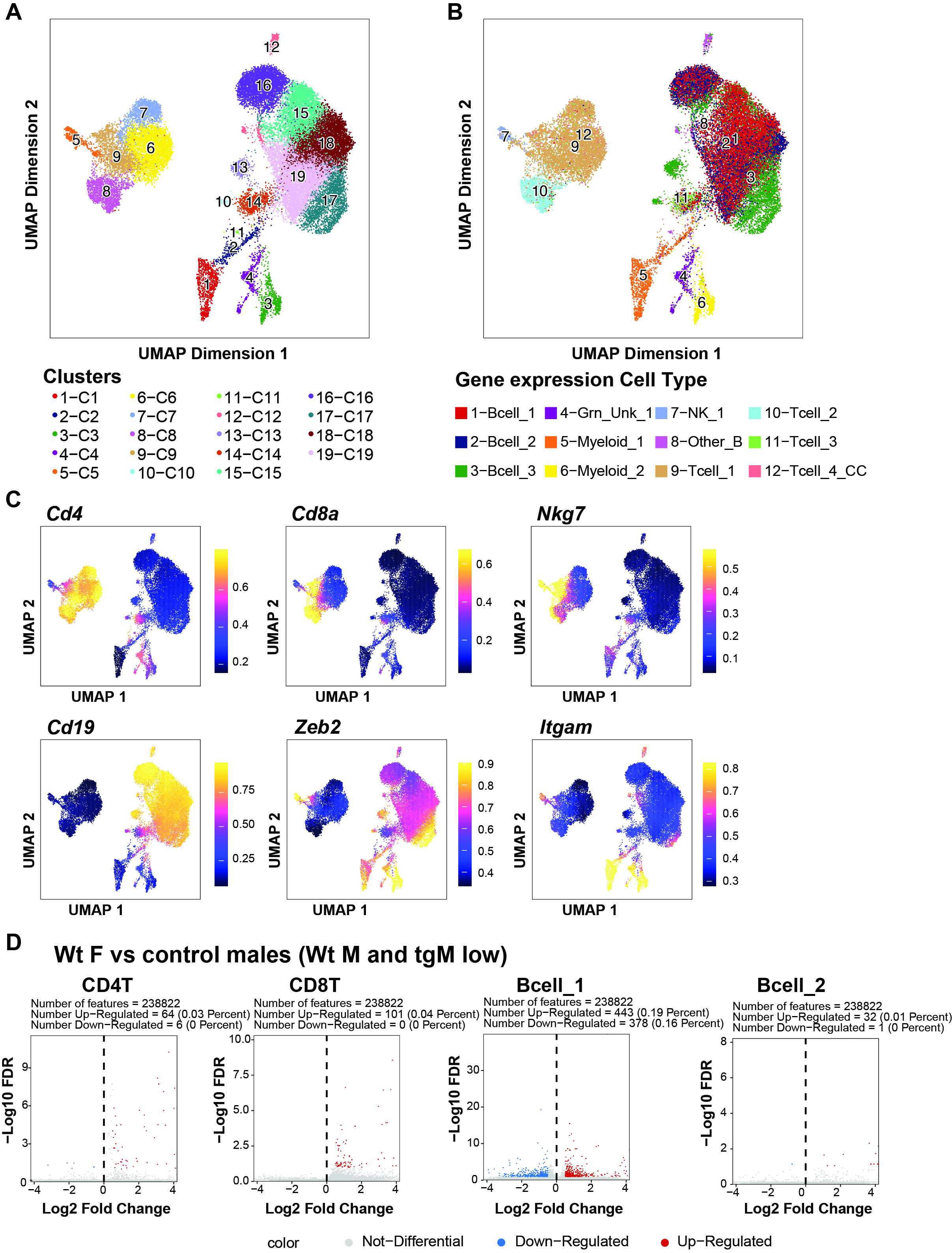

**Supplementary Figure 4: Splenic CD45+ hematopoietic cells single-cell ATAC from pristane-induced SLE mice of the SJL/J strain:** (A) Original single-cell ATAC clusters and (B) matched single cell gene expression-determined cell identities displayed on the single cell ATAC UMAP. (C) Localization of defining markers, calculated by imputation from ATAC data, used to determine cell type identity. Imputation scale Log2(NormCounts+1) (D) Pairwise comparison metrics of differential peaks between WT female (positive disease control) and the low disease male control groups (WT Male and tgXist Male low disease) across the four main cellular subsets shown for features FDR ≦ 0.1 and Log2FC ≧ 0.5. Pristane-treated mouse groups shown: tgXist male disease high (n=4), tgXist male disease low (n=4), wild-type male (n=3), and wild-type female (n=2). Significance was calculated using the Wilcoxon Rank Sum test.

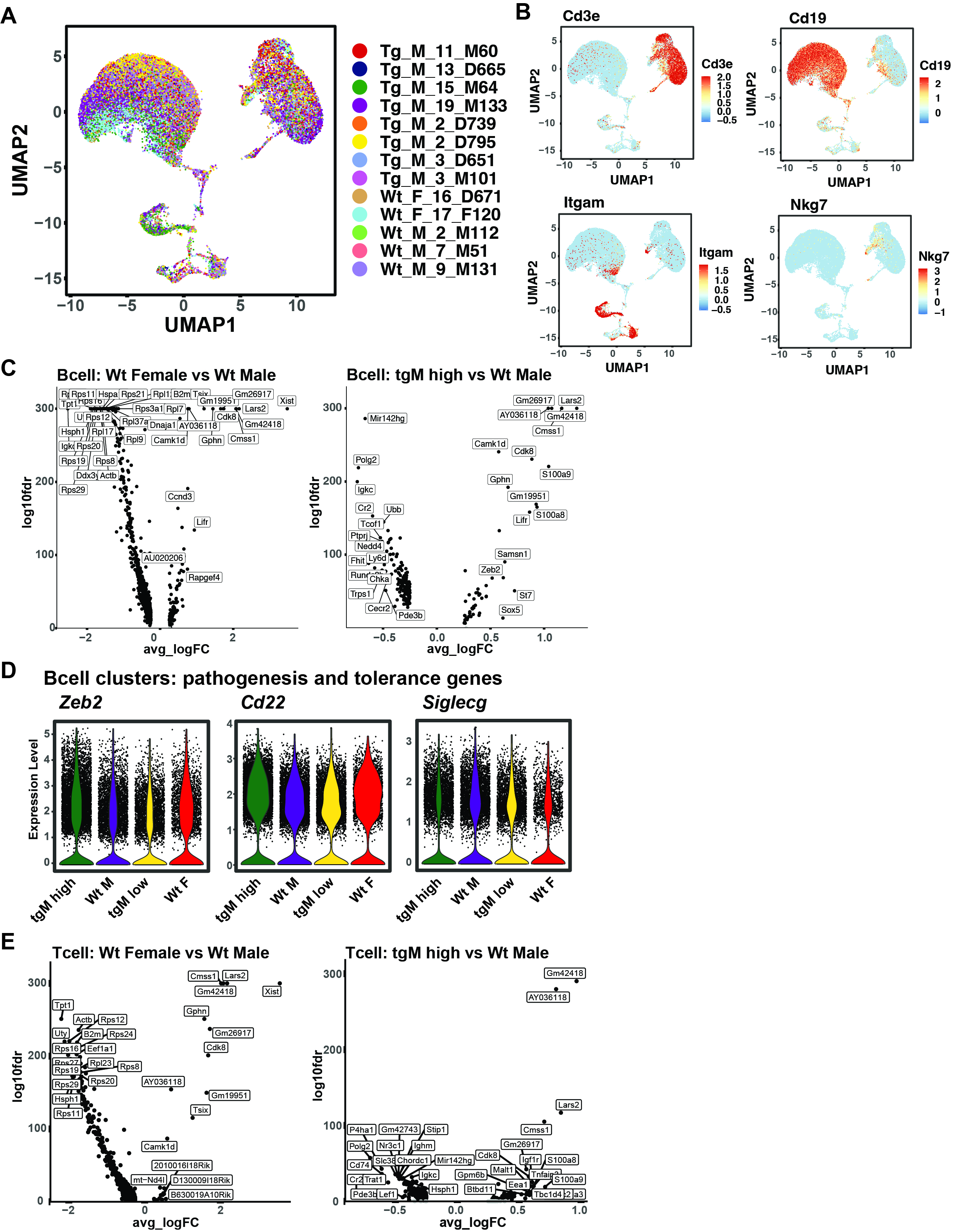

**Supplementary Figure 5: Splenic CD45+ hematopoietic cells single-cell gene expression from pristane-induced SLE mice of the SJL/J strain:** (A) Individual mice displayed on the single cell gene expression clustering UMAP. Individual mouse labels displayed as: transgenic status_sex_total disease damage score_mouse colony ID. (B) Localization of defining expression markers used to determine immune cell type identities of UMAP clusters. Volcano plots of differentially expressed (C) B cell cluster genes comparing tgXist Male high disease and WT Female to WT Male. (D) Representative violin plots of B-cell marker genes from the combined B cell clusters and (E) T cell cluster genes comparing tgXist Male high disease and WT Female to WT Male. Pristane-treated mouse groups shown: tgXist male disease high (n=4), tgXist male disease low (n=4), wild-type male (n=3), and wild-type female (n=2). Significance was calculated

using the Wilcoxon Rank Sum test.

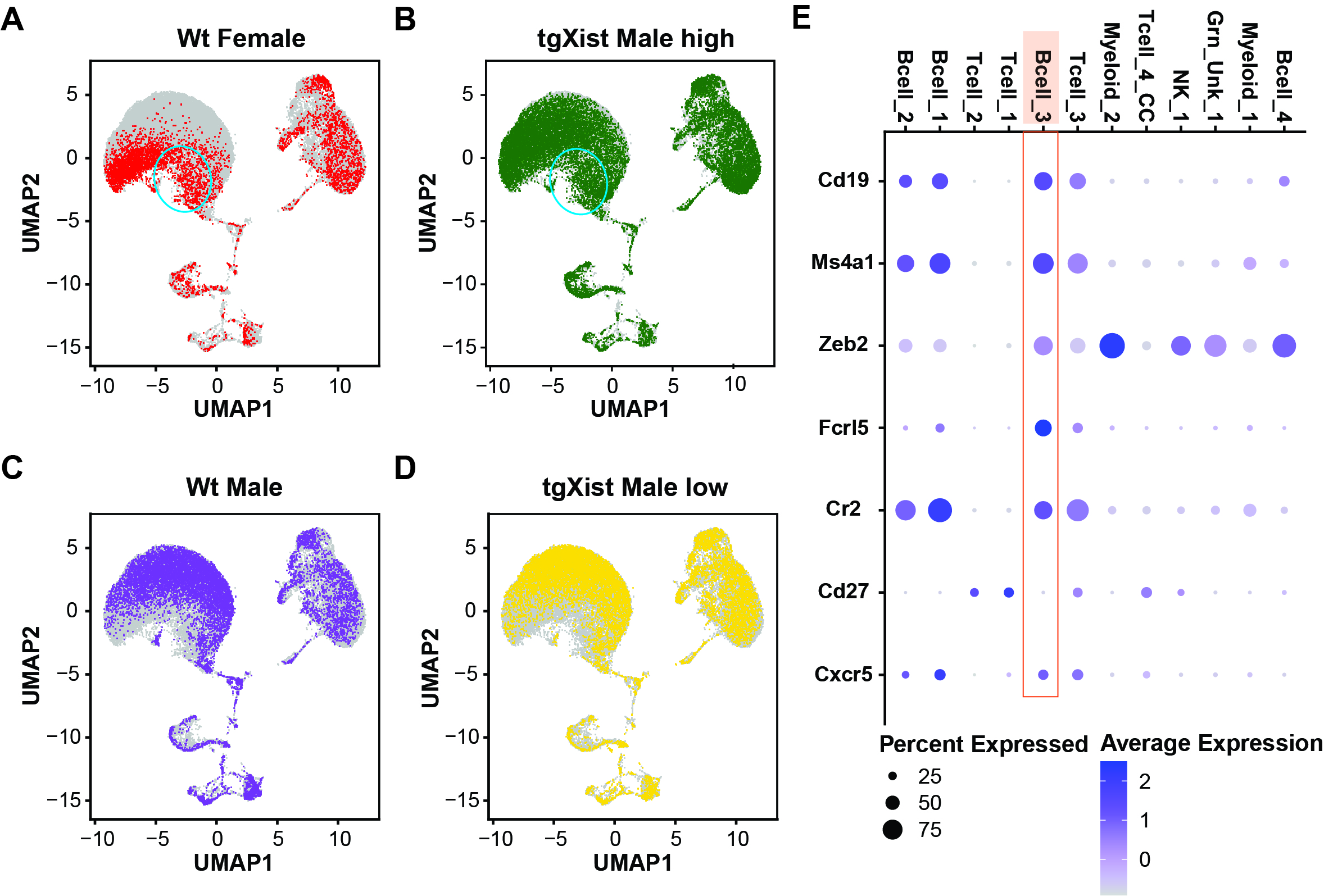

**Supplementary Table 6: Distribution of pristane-treated mouse cohorts and atypical B cells.** UMAP distribution of cells from (A) Wt female, (B) tgXist Male high disease, (C) Wt Male and (D) tgXist Male low diseases. Blue circles indicate shared overlapping regions in Wt female and tgXist Male high disease pristane-treated mouse groups corresponding to Bcell_3. (E) Dot plot of atypical B-cell marker expression in gene expression clusters. Pristane-treated mouse groups shown: tgXist male disease high (n=4), tgXist male disease low (n=4), wild-type male (n=3), and wild-type female (n=2)

**Supplementary Tables**

Numbers indicate Sheets in Excel files

**Supplementary Table 1**: **XIST Bibliomics**. Identification of known autoantigens in the XIST RNP and their associated autoimmune diseases and publications. 1) XIST-associated proteins were drawn from published and validated XIST ChIRP-MS datasets. 2) XIST-associated proteins classified as reactive of enriched from the XIST antigen array that were not included in the prior ChIRP-MS lists of high stringency validated XIST RNP proteins.

**Supplementary Table 2: Pristane-induced SLE mouse study controls and design.** Study design and appropriate matched control comparisons of each tgXist and wild-type mouse treatment cohort in the 1) C57BL/6J and 2) SJL/J backgrounds.

**Supplementary Table 3**: **Pristane-induced SLE antigen array**. Raw Mean Fluorescence intensity (MFI) values and mouse serum sample information of pristane-induced SLE studies to SLE autoantigens in wild-type and tgXist 1), 2) C57/BL6J and 3), 4) SJL/J mice.

**Supplementary Table 4**: **XIST Antigen Array in patients**. 1) Description and IDs of antigens used in the array to evaluate patient sera reactivity. Rows are labeled with “Gene name_Antigen name”, columns describe association (assay control, clinical disease autoantibody, XIST ChIRP, or Exploratory XIST), gene names, IDs in public databases, control or test status, multiplicity of targets for each fragment (also indicated as a “*” in the row labels as “Gene name*_Antigen name”), and fragment sequence for each antigen. 2) Sample disease and biological sex information. 3) Raw Mean Fluorescence intensity (MFI) values of patient serum reactivity to antigens.

**Supplementary Table 5**: **Reactive Antigens in Autoimmune Disease Patients**. List of all reactive antigens in the three autoimmune disease patient cohorts evaluated in this study and known status as autoantigens. XIST RNP complex proteins in blue text. Highlighted antigens correspond to the published high confidence XIST RNP complex proteins^14^. “Autoantigen Record” indicates status of the antigen as a known disease control (Control), present in **Supplementary Table 1** **XIST Bibliomics** publications (Known), or not present in included Bibliomics publications (Unknown). Type of significantly reactive autoimmune disease cohorts indicated in “Array Reactivity”.

**Supplementary Table 6**: **XIST Antigen Array in SJL/J mice**. 1) Description and IDs of antigens used in the array to evaluate mouse sera reactivity. Rows are labeled with “Gene name_Antigen name”, columns describe association (assay control, clinical disease autoantibody, XIST ChIRP, or Exploratory Xist), gene names, IDs in public databases, control or test status, multiplicity of targets for each fragment (also indicated as a “*” in the row labels as “Gene name*_Antigen name”), and fragment sequence for each antigen. 2) Sample treatment, timepoint and sex information. 3) Raw Mean Fluorescence intensity (MFI) values of mouse sera reactivity to antigens.

**Supplementary Table 7**: **Enriched Antigens in SLE**. List of all antigens enriched with outlier-trained MAD difference > 0 and raw MAD difference >2.5-fold MAD scores compared to general population (Described in Methods). XIST RNP complex proteins in blue text. Highlighted antigens correspond to the published high confidence XIST RNP complex proteins^14^. “Autoantigen Record” indicates status of the antigen as a known disease control (Control), present in **Supplementary Table 1** **XIST Bibliomics** publications (Known), or not present in included Bibliomics publications (Unknown). Corresponding mouse cohort overlap indicated in “Mouse Model Reactivity”.

**Supplementary Table 8**: **Primers**. Sequences for genotyping and qRT-PCR primers used in the study.

**Supplementary Table 9**: **Pristane-induced SLE in SJL/J Mice Histology**. Complete full body organ histology scores for all SJL/J mice in the study. Due to pandemic difficulties, a subset of mice were “affected by processing” that rendered the isolated kidney unusable for downstream analysis (indicated as “Y” in the column and “?” under glomerulonephritis) and some mice were missing splenic and lymph node scores. Only mice with all organ scores (in bold text) were included in the final **Figure 3** Total Organ Pathology statistical analysis.

**Supplementary Table 10**: **Single Cell Multiome Quality Check**. Reads and quality score metrics for Single Cell Multiome ATAC + Gene Expression sequencing libraries 1) ATAC-seq and 2) Gene Expression.
